# Two Ascending Thermosensory Pathways from the Lateral Parabrachial Nucleus That Mediate Behavioral and Autonomous Thermoregulation

**DOI:** 10.1101/2023.04.10.536301

**Authors:** Takaki Yahiro, Naoya Kataoka, Kazuhiro Nakamura

**Affiliations:** Department of Integrative Physiology, Nagoya University Graduate School of Medicine, Nagoya 466-8550, Japan; Nagoya University Institute for Advanced Research, Nagoya 464-8601, Japan

## Abstract

Thermoregulatory behavior in homeothermic animals is an innate behavior to defend body core temperature from environmental thermal challenges in coordination with autonomous thermoregulatory responses. In contrast to the progress in understanding the central mechanisms of autonomous thermoregulation, those of behavioral thermoregulation remain poorly understood. We have previously shown that the lateral parabrachial nucleus (LPB) mediates cutaneous thermosensory afferent signaling for thermoregulation. To understand the thermosensory neural network for behavioral thermoregulation, in the present study, we investigated the roles of ascending thermosensory pathways from the LPB in avoidance behavior from innocuous heat and cold in rats. Neuronal tracing revealed two segregated groups of LPB neurons projecting to the median preoptic nucleus (MnPO), a thermoregulatory center (LPB→MnPO neurons), and those projecting to the central amygdaloid nucleus (CeA), a limbic emotion center (LPB→CeA neurons). While LPB→MnPO neurons include separate subgroups activated by heat or cold exposure of rats, LPB→CeA neurons were only activated by cold exposure. By selectively inhibiting LPB→MnPO or LPB→CeA neurons using tetanus toxin light chain or chemogenetic or optogenetic techniques, we found that LPB→MnPO transmission mediates heat avoidance, whereas LPB→CeA transmission contributes to cold avoidance. *In vivo* electrophysiological experiments showed that skin cooling-evoked thermogenesis in brown adipose tissue requires not only LPB→MnPO neurons but also LPB→CeA neurons, providing a novel insight into the central mechanism of autonomous thermoregulation. Our findings reveal an important framework of central thermosensory afferent pathways to coordinate behavioral and autonomous thermoregulation and to generate the emotions of thermal comfort and discomfort that drive thermoregulatory behavior.

## Introduction

Homeothermic animals employ voluntary behavioral and involuntary autonomous thermoregulation in a coordinated manner to maintain body core temperature (*T*_core_) within a narrow range against changes in environmental temperature. Thermoregulatory behaviors, such as thermal comfort selection (*i.e.*, avoidance of hot and cold temperatures), postural extension, grooming, and locomotion, enhance the effectiveness of autonomous thermoregulatory responses, such as sympathetic thermogenesis in brown adipose tissue (BAT), shivering thermogenesis in skeletal muscles, and cutaneous vasomotion. Despite the remarkable progress in understanding the central circuit mechanisms of autonomous thermoregulation (Morrison and Nakamura, 2019; Nakamura et al., 2022a), those of behavioral thermoregulation are still under investigation.

Changes in environmental temperature are sensed by cutaneous warm and cold receptors and primary somatosensory fibers separately transmit the warm and cold sensory signals to lamina Ⅰ neurons in the spinal and medullary dorsal horns, which then transmit the signals to the thalamus and to the lateral parabrachial nucleus (LPB) in the pons (Hylden et al., 1989; Li et al., 2006). The spinothalamic pathway delivers the sensory signals to the primary somatosensory cortex for the perception and discrimination of skin temperature (Craig et al., 1994; Craig, 2002). On the other hand, the spinoparabrachial pathway extends thermosensory signaling to the preoptic area (POA), a thermoregulatory center, to elicit immediate autonomous thermoregulatory responses to the changes in environmental temperature (Nakamura and Morrison, 2008, 2010). We have shown that the LPB, but not the thalamus, also mediates thermosensory signaling for avoidance of innocuous heat and cold (Yahiro et al., 2017). However, despite recent efforts (Norris et al., 2021; Jung et al., 2022), thermosensory neural pathways ascending from the LPB for behavioral thermoregulation are largely unknown.

In the LPB, two major groups of neurons expressing different transcription factors, forkhead box protein P2 (FoxP2) and LIM homeobox transcription factor 1β (Lmx1b), show segregated distributions (Miller et al., 2012). FoxP2-expressing LPB neurons project numerous axons to the POA and other hypothalamic regions (Huang et al., 2021a). In contrast, the Lmx1b-expressing group includes calcitonin gene-related peptide (CGRP)-expressing neurons that densely innervate the central amygdaloid nucleus (CeA) and other limbic regions (Huang et al., 2021b). It is hypothesized that these two projection groups transmit different thermosensory signals to the forebrain sites to drive coordinated behavioral and autonomous thermoregulatory responses. However, whether the POA is involved in behavioral thermoregulation has been controversial (Almeida et al., 2015). Whereas lesioning the POA does not affect thermal preference or operant thermoregulatory behaviors (Lipton and Hicks, 1968; Carlisle, 1969; Almeida et al., 2006), the POA is involved in body extension of rodents in a warm environment (Carlisle, 1969; Satinoff and Rutstein, 1970; Yu et al., 2016; Norris et al., 2021). Also, activation of PACAP/BDNF-positive POA neurons elicits cold-seeking behavior and inhibits nest building (Tan et al., 2016). Although these findings implicate the POA in behavioral thermoregulation, the importance of LPB→POA thermosensory transmission in behavioral thermoregulation remains unclear.

The LPB→CeA pathway mediates nociceptive signaling to generate a fear memory that induces aversive behavior to pain (Bernard and Besson, 1990; Gauriau and Bernard, 2002; Han et al., 2015; Sato et al., 2015). However, whether the LPB→CeA pathway is involved in thermoregulatory avoidance of innocuous hot and cold is unknown. Because thermoregulatory behaviors are motivated by thermally evoked comfort and discomfort, it would be relevant to investigate whether emotion-related corticolimbic circuits, including the amygdala, are involved in behavioral thermoregulation.

In the present study, we investigated the roles of LPB neurons projecting to the median preoptic nucleus (MnPO) and those projecting to the CeA in behavioral thermoregulation and in coordination with autonomous thermoregulation in rats. We performed functional neuroanatomy to examine the distribution of collateral axons and the thermal responsiveness of the two projection neurons. Using tetanus toxin light chain (TeTxLC), designer receptors exclusively activated by designer drugs (DREADDs), and inhibitory optogenetic channels, we chronically or acutely inhibit LPB→MnPO or LPB→CeA transmission selectively, and performed two-floor innocuous thermal plate preference tests (TPPTs) to assess heat and cold avoidance behaviors and *in vivo* physiological recordings of autonomous thermoregulatory responses.

## Materials and Methods

### Animals

Male Wistar ST rats (280–400g; 8–12 weeks old) were purchased from Japan SLC (Hamamatsu, Japan) and housed with *ad libitum* access to food and water in a room air-conditioned at 25 ± 2°C with a standard 12-h light/dark cycle (light: 7:00–19:00, dark: 19:00– 7:00) until they were used for surgery or experiments. All procedures conform to the guidelines of animal care by the Division of Experimental Animals, Nagoya University Graduate School of Medicine, and were approved by the Nagoya University Animal Care and Use Committee (approval No. M220097-002).

### Adeno-associated virus (AAV) vectors

Recombinant AAVs used in the present study except AAVrg-pgk-Cre were serotype 1 or 8 (see below). AAV-CMV-palGFP, which was used to transduce LPB neurons with a membrane-targeted form of green fluorescent protein (palGFP), was described in our earlier study (Kataoka et al., 2014). AAVs for Cre-dependent expression of iChloC-mCherry (AAV-EF1α-DIO-iChloC-mCherry) and palGFP (AAV-EF1α-DIO-palGFP) were also described previously (Kataoka et al., 2020). An AAV retrograde (AAVrg) for Cre expression (AAVrg-pgk-Cre) was purchased from Addgene (Addgene, #24593, donated by Patrick Aebischer).

To produce AAVs for Cre-dependent expression of TeTxLC and EYFP (AAV-EF1α-DIO-TeTxLC-T2A-EYFP) and hM4Di^nrxn^ and mCherry (AAV-EF1α-DIO-mCherry-T2A-hM4Di^nrxn^), we inserted a DNA cassette encoding TeTxLC-T2A-EYFP (VectorBuilder) or mCherry-T2A-hM4Di^nrxn^ (Addgene, #52523, donated by Scott Sternson) into the plasmid, pAAV2-Ef1α-DIO-MCS, respectively. The produced plasmids, pAAV2-EF1α-DIO-TeTxLC-T2A-EYFP and pAAV2-Ef1α-DIO-mCherry-T2A-hM4Di^nrxn^, were used for the production and purification of the AAVs according to our methods (Kataoka et al., 2020; Takahashi et al., 2021).

The titrations of the AAVs used were 3.5 × 10^11^ GC/ml for AAV2/1-CMV-palGFP, 1.6 × 10^14^ GC/ml for AAV2/8-EF1α-DIO-iChloC-mCherry, 1.1 × 10^14^ GC/ml for AAV2/8-EF1α-DIO-palGFP, 9.5 × 10^12^ GC/ml for AAVrg-pgk-Cre, 6.0 × 10^12^ GC/ml for AAV2/8-EF1α-DIO-TeTxLC-T2A-EYFP, and 1.7 × 10^14^ GC/ml for AAV2/8-EF1α-DIO-mCherry-T2A-hM4Di^nrxn^.

### Surgery

Rats were anesthetized by a subcutaneous injection with a combination anesthetic [medetomidine hydrochloride (0.15 mg/kg), midazolam (2.0 mg/kg), and butorphanol tartrate (2.5 mg/kg)] following gas anesthesia with 3% isoflurane. The depth of anesthesia was carefully checked by the absence of hind limb withdrawal to a foot pinch and/or by the absence of eye blinking in response to a gentle touch to the cornea before the rats were positioned in a stereotaxic apparatus. During surgery, the appropriate depth of anesthesia was maintained by ensuring that no sign of pain or discomfort was observed.

A glass micropipette filled with a solution containing AAV or cholera toxin b-subunit (CTb) conjugated with Alexa Fluor 488 (Alexa488) or Alexa594 (1 mg/ml; C22841 and C22842, Thermo Fisher Scientific) was perpendicularly inserted into the LPB, MnPO or CeA. The solution was pressure-ejected using a Picospritzer III (Parker, Hollis, NH), and then, the micropipette was left in place for 5 min before being withdrawn. AAV injections were targeted at the following coordinates: LPB (9.0 mm caudal to bregma, 2.3 mm left and right to the midline, and 6.0 mm ventral to the brain surface; 200 nl/site), MnPO (on bregma, on the midline, and 7.4 and 7.7 mm ventral to the brain surface; 150 nl/site) and CeA (2.0 to 3.0 mm caudal to bregma, 4.7 mm left and right to the midline, and 7.6 mm ventral to the brain surface; 150 nl/site). CTb injection was made into the MnPO (coordinates as above; 150 nl/site) and bilaterally into the CeA (coordinates as above; 200 nl/site).

For intracranial drug nanoinjections in DREADD experiments, a single sterile guide cannula (ID = 0.24 mm, OD = 0.46 mm; C315G; Plastic One, Roanoke, VA) was inserted to target the MnPO (coordinates: 0.1 mm rostral to bregma, 1.4 mm left to the midline, and 6.6 mm ventral to the brain surface, tilted 10° laterally) or a paired sterile bilateral stainless guide cannula (ID = 0.24 mm, OD = 0.46 mm, inter-cannula distance = 1.6 mm; C235G-1.6/SPC, Plastics One, Roanoke, VA) was perpendicularly inserted to target the DMH (coordinates: 2.8 mm caudal to bregma, 0.8 mm left and right to the midline, and 6.8 mm ventral to the brain surface) after AAV injections. The guide cannulae were then anchored with dental cement to stainless steel screws attached to the skull. Dummy cannulae cut to the exact length of the guide cannulae were inserted into the guide cannulae to avoid clogging. Internal cannulae for injection with the thickness to fit the guide cannulae were cut to be long enough to allow the injector tip to protrude 1.0 mm below the tip of the guide cannulae.

For optogenetic experiments, fiber optic cannulae with a ceramic ferrule (200 μm core diameter, 0.39 NA; R-FOC-BL200C-39NA, RWD, San Diego, CA) were bilaterally inserted to target 0.5 mm dorsal to the CeA (coordinates: 2.5 mm caudal to bregma, 4.7 mm left and right to the midline, and 7.1 mm ventral to the brain surface) or ventromedial hypothalamic nucleus (VMH; coordinates: 2.8 mm caudal to bregma, 2.1 mm left and right to the midline, and 8.9 mm ventral to the brain surface, tilted 10° laterally).

For telemetric monitoring of *T*_core_, the anesthetized rats were also implanted with a battery-operated telemetric transmitter (TA-F40, Data Science International, St. Paul, MN) after the above procedures. The abdominal wall was incised and the telemetric transmitter was placed in the abdominal cavity.

At the end of surgery, the incised skin and abdominal wall were closed with sutures or wound clips and the surgical wound was treated with iodine. Ampicillin sodium solution (0.2 ml, 125 mg/ml) was injected into the left femoral muscle to prevent postoperative infection. Atipamezole solution (0.1 ml, 0.5 mg/ml) was injected subcutaneously to promote arousal. The rats were caged individually for at least one week to recover from the surgery with *ad libitum* access to food and water in a room air-conditioned at 24 ± 2°C under regular health checks.

### Neural tracing and cold and heat exposure

Rats injected with AAV-CMV-palGFP or CTb were transcardially perfused as described below, > 1 week after injection. For retrograde tracing with CTb combined with Fos study, CTb-injected, individually caged rats were exposed to 4°C, 25°C or 36°C for 2 h in a climate chamber as described (Nakamura et al., 2022b), > 1 week after CTb injection. Immediately after the exposure, the animals were subjected to transcardial perfusion below.

### TPPT

The apparatus consisted of two black-painted thermal plates (19 cm × 30 cm) placed side by side, surrounded by black acrylic walls (40 cm high) with a slit for passage between the two plates (Fig. 1F). The apparatus was placed in a climate chamber air-conditioned at 25°C ± 0.5°C. A controller (Intercross, Tokyo, Japan) regulated the temperature of each plate within an error of ± 0.1°C using Peltier devices. For experiments with rats, one plate was set at 28°C and the other at 15°C or 39°C, as we performed previously (Yahiro et al., 2017). The temperature settings of the plates were randomly assigned, but not side-fixed. Immediately after a rat was placed on the plates, its behavior was monitored and recorded with a camera (C920 Pro HD Webcam, Logitech) for 20 or 30 min, and the time spent on each plate, the timing of transitions between plates, and the distance traveled were analyzed with the video-tracking software, ANY-maze (Stoelthing, Wood Dale, IL). The percentage of time spent on each plate during the test period was calculated. During testing, *T*_core_ was monitored by a telemetry system (Ponemah, Data Science International, St. Paul, MN).

**Figure 1.**
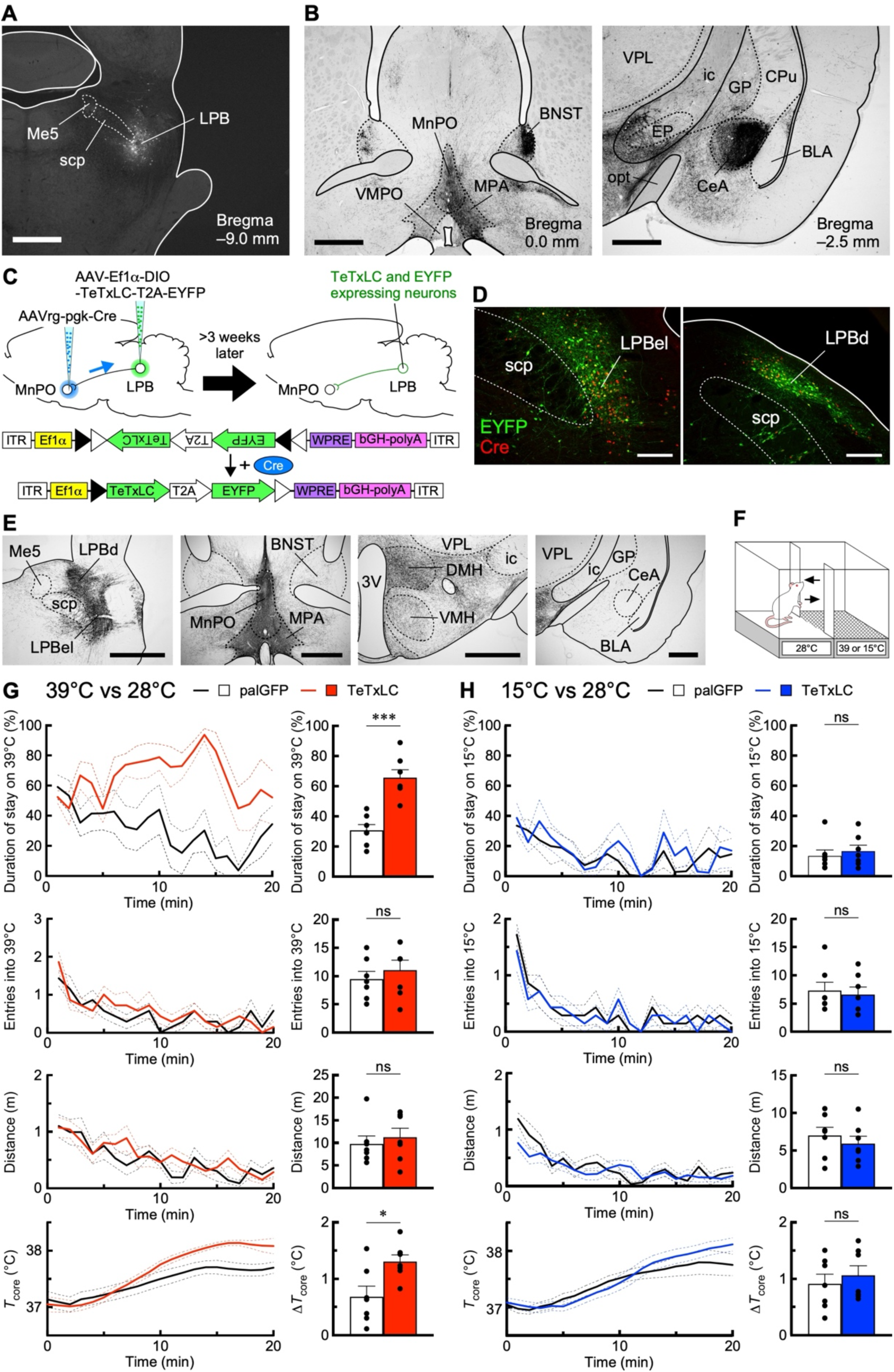
TeTxLC-mediated chronic suppression of LPB→MnPO neurons abolishes heat avoidance, but not cold avoidance. **A and B**, Non-selective AAV-transduction of LPB neurons with palGFP (A) visualized their axons densely distributed in the MnPO and the CeA (B). Scale bars, 1.0 mm. BLA, basolateral amygdaloid nucleus; BNST, bed nucleus of the stria terminalis; CPu, caudate putamen; EP, entopeduncular nucleus; GP, globus pallidus; ic, internal capsule; Me5, mesencephalic trigeminal nucleus; MPA, medial preoptic area; scp, superior cerebellar peduncle; VMPO, ventromedial preoptic nucleus; VPL, ventral posterolateral thalamic nucleus. **C**, Selective transduction of LPB→MnPO neurons with EYFP-T2A-TeTxLC using AAV and AAVrg. bGH-polyA, bovine growth hormone polyadenylation signal; Ef1α, elongation factor 1-α promoter; ITR, inverted terminal repeat; WPRE, woodchuck hepatitis virus posttranscriptional regulatory element. **D**, Cre- and/or EYFP-immunoreactive neurons in the LPB. Scale bars, 500 μm. **E**, Selective transduction of LPB→MnPO neurons with palGFP visualized distributions of axons of LPB→MnPO neurons. Scale bars, 1 mm. 3V, third ventricle. **F**, Thermal plate preference test. **G and H**, Effects of TeTxLC-expression in LPB→MnPO neurons on heat (G) and cold (H) avoidance. Control rats expressed palGFP instead of EYFP-T2A-TeTxLC. Time-course changes (left, every 1 min) in % duration of stay on 39°C or 15°C plate, times of entry into 39°C or 15°C plate, distance traveled, and *T*_core_. Right graphs show % duration of stay on 39°C or 15°C plate, total times of entry into 39°C or 15°C plate, total distance traveled, and change in *T*_core_ (Δ*T*_core_, difference between value at time 0 and peak) for the 20-min test period, and the data were analyzed by unpaired *t*-tests (*n* = 7 per group; (G): duration of stay, *t*_12_ = 5.39; entries, *t*_12_ = 0.70; distance, *t*_12_ = 0.56; Δ*T*_core_, *t*_12_ = 2.80; (H): duration of stay, *t*_12_ = 0.57; entries, *t*_12_ = 0.36; distance, *t*_12_ = 0.74; Δ*T*_core_, *t*_12_ = 0.63). \**P* < 0.05; \*\*\**P* < 0.001; ns, not significant. Error bars indicate SEM.

In DREADD experiments, the rats that had undergone intracranial cannulation received bilateral nanoinjections of either saline or C21 into the MnPO or DMH through the cannulae prior to a TPPT. An internal cannula was connected to a Teflon tubing, and the inside of the cannula and tubing was filled with pyrogen-free 0.9% saline (Otsuka, Tokyo, Japan) or C21 (200 μg/ml, dissolved in saline; SML2392, Sigma-Aldrich). A Hamilton syringe (10 μl) filled with mineral oil was connected to the other end of the Teflon tubing and placed in a manually operated syringe manipulator (Narishige, Tokyo, Japan). The dummy cannulae inserted into the guide cannulae on the skull were gently removed, and the internal cannula was inserted into one of the guide cannulae. Then, the saline or C21 solution (50 nl/side) was slowly ejected through the cannula into the MnPO or DMH using the syringe manipulator. The injection volume was visually confirmed by the movement of the aqua–oil interface along the Teflon tubing. For bilateral injections into the DMH, immediately after injection on one side, another injection was made on the other side. Five minutes after completion of the bilateral nanoinjections, the rats were subjected to a 30-min TPPT.

For optogenetic experiments, the fiber optic cannulae implanted on the skull were connected with ferrule patch cables (M81L01, Thorlabs). The light source was a diode-pumped 445 nm blue laser (LuxX 445-100, Omicron, Dudenhofen, Germany) controlled by a function generator (AFG1022, Tektronix). The power output was measured at the fiber tip with a light meter beforehand and preset at 8 mW (when the laser was activated in a continuous mode). The target sites were illuminated with 50-ms light pulses at 10 Hz immediately after the rats were placed on the thermal plates, and illumination was continued for the 20 min of a TPPT period.

### *In vivo* electrophysiology

*In vivo* physiological recordings followed our established procedure (Nakamura and Morrison, 2007, 2011). AAV-injected rats were anesthetized intravenously with urethane (0.8 g/kg) and α-chloralose (80 mg/kg) after cannulation of a femoral artery, a femoral vein, and the trachea under anesthesia with 3% isoflurane in 100% O_2_. The arterial cannula was attached to a pressure transducer to record arterial pressure and heart rate (HR). Body core temperature (*T*_core_) was monitored by a copper-constantan thermocouple (Physitemp) inserted into the rectum. The trunk was shaved and wrapped with a plastic water jacket, and skin temperature (*T*_skin_) was monitored from a thermocouple taped onto the skin 1 cm caudal to the xiphoid process. The water jacket was perfused with warm or cold water to maintain the *T*_core_ between 36.0 and 38.0°C. The rats were placed in a stereotaxic apparatus, artificially ventilated with 100% O_2_ through the tracheal cannula, and paralyzed with D-tubocurarine (0.6 mg, i.v., initial dose, supplemented with 0.3 mg/h) to stabilize BAT sympathetic nerve recording by preventing respiration-related movements. Mixed expired CO_2_ was monitored through the tracheal cannula using a capnometer to provide an index of changes in whole-body metabolism and was maintained between 3.5 and 4.5% under basal conditions. BAT temperature (*T*_BAT_) was measured from the left interscapular BAT pad with a needle-type copper-constantan thermocouple (0.33 mm diameter; Physitemp), and postganglionic BAT sympathetic nerve activity (SNA) was recorded from the central cut end of a nerve bundle isolated from the right interscapular BAT pad. BAT SNA was amplified (× 5,000 to 10,000) and filtered (1 to 300 Hz) using a CyberAmp 380 amplifier (Molecular Devices). All the physiological variables were digitized and recorded to a personal computer using Spike2 software (version 7.10; CED, Cambridge, UK). BAT SNA amplitudes were quantified using Spike2 in sequential 4-s bins as the square root of the total power (root mean square) in the 0- to 20-Hz band of the autospectra of each 4-s segment of the BAT SNA traces. The “power/4 s” traces (Fig. 2A, B, D and E and Fig. 7A, B, D and E) were used for quantification and statistical analyses of changes in BAT SNA.

**Figure 2.**
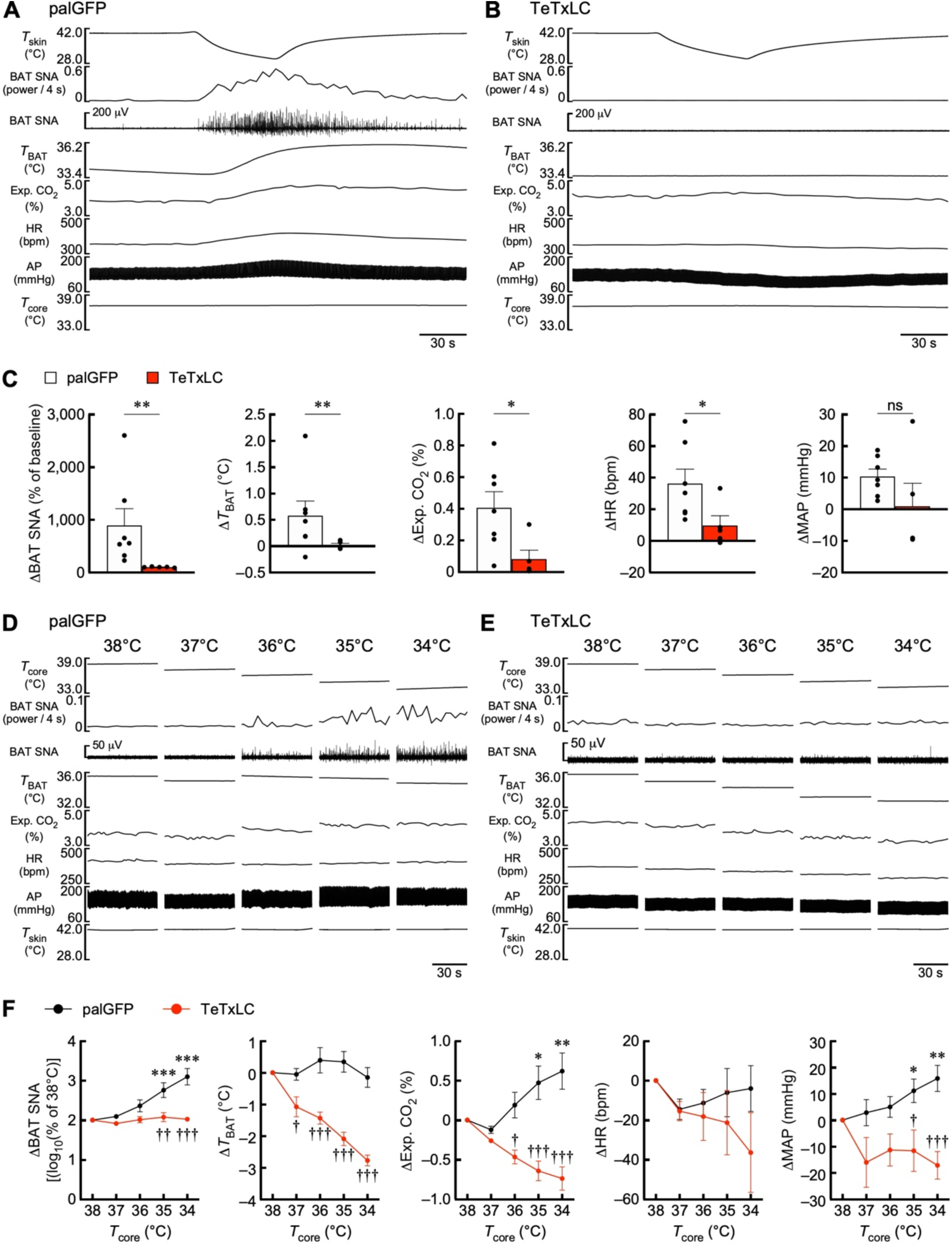
LPB→MnPO neurons mediate thermosensory signaling for BAT thermogenic and cardiovascular responses to skin cooling and body core cooling. **A and B**, Effects of palGFP- and TeTxLC-expression in LPB→MnPO neurons on BAT thermogenic and cardiovascular responses to skin cooling. AP, arterial pressure; Exp. CO_2_, expired CO_2_. **C**, Skin cooling-evoked changes in physiological variables, which were analyzed by Mann-Whitney U tests (palGFP: *n* = 7, TeTxLC: *n* = 5; ΔBAT SNA: *U* = 0; Δ*T*_BAT_: *U* = 5; ΔExp. CO_2_: *U* = 4; ΔHR: *U* =; 4; ΔMAP: *U* = 9). **D and E**, Effects of palGFP- and TeTxLC-expression in LPB→MnPO neurons on BAT thermogenic and cardiovascular responses to reduction of *T*_core_. **F**, Body core cooling-evoked changes in BAT SNA, *T*_BAT_, Exp. CO_2_, MAP, and HR were compared to the values at 38°C. Data from control (palGFP) rats (*n* = 7) were analyzed by repeated measures one-way ANOVA followed by Bonferroni’s post hoc test (ΔBAT SNA: *F*_4, 24_ = 21.53, *P* < 0.001; Δ*T*_BAT_: *F*_4, 24_ = 1.74, *P* = 0.175; ΔExp. CO_2_: *F*_4, 24_ = 7.91, *P* < 0.001; ΔHR: *F*_4, 24_ = 0.99, *P* = 0.434; ΔMAP: *F*_4, 24_ = 5.39, *P* = 0.003). \**P* < 0.05; \*\**P* < 0.01; \*\*\**P* < 0.001 (vs 38°C). Differences in cooling-evoked changes in the physiological variables between palGFP and TeTxLc groups were analyzed by repeated measures two-way ANOVA followed by Bonferroni’s post hoc test (palGFP, *n* = 7; TeTxLC, *n* = 5; ΔBAT SNA: group: *F*_1, 10_ = 10.84, *P* = 0.008, *T*_core_: *F*_4, 40_ = 15.34, *P* < 0.001, interaction: *F*_4, 40_ = 11.19, *P* < 0.001; Δ*T*_BAT_: group: *F*_1, 10_ = 31.39, *P* < 0.001, *T*_core_: *F*_4, 40_ = 14.60, *P* < 0.001, interaction: *F*_4, 40_ = 14.88, *P* < 0.001; ΔExp. CO_2_: group: *F*_1, 10_ = 16.87, *P* = 0.002, *T*_core_: *F*_4, 40_ = 1.02, *P* = 0.410, interaction: *F*_4, 40_ = 16.61, *P* < 0.001; ΔHR: group: *F*_1, 10_ = 0.98, *P* = 0.346, *T*_core_: *F*_4, 40_ = 1.97, *P* = 0.117, interaction: *F*_4, 40_ = 1.54, *P* = 0.208; ΔMAP: group: *F*_1, 10_ = 10.12, *P* < 0.001, *T*_core_: *F*_4, 40_ = 1.19, *P* = 0.332, interaction: *F*_4, 40_ = 5.44, *P* = 0.001). ^†^*P* < 0.05; ^††^*P* < 0.01; ^†††^*P* < 0.001 (vs palGFP). Error bars indicate SEM.

**Figure 7.**
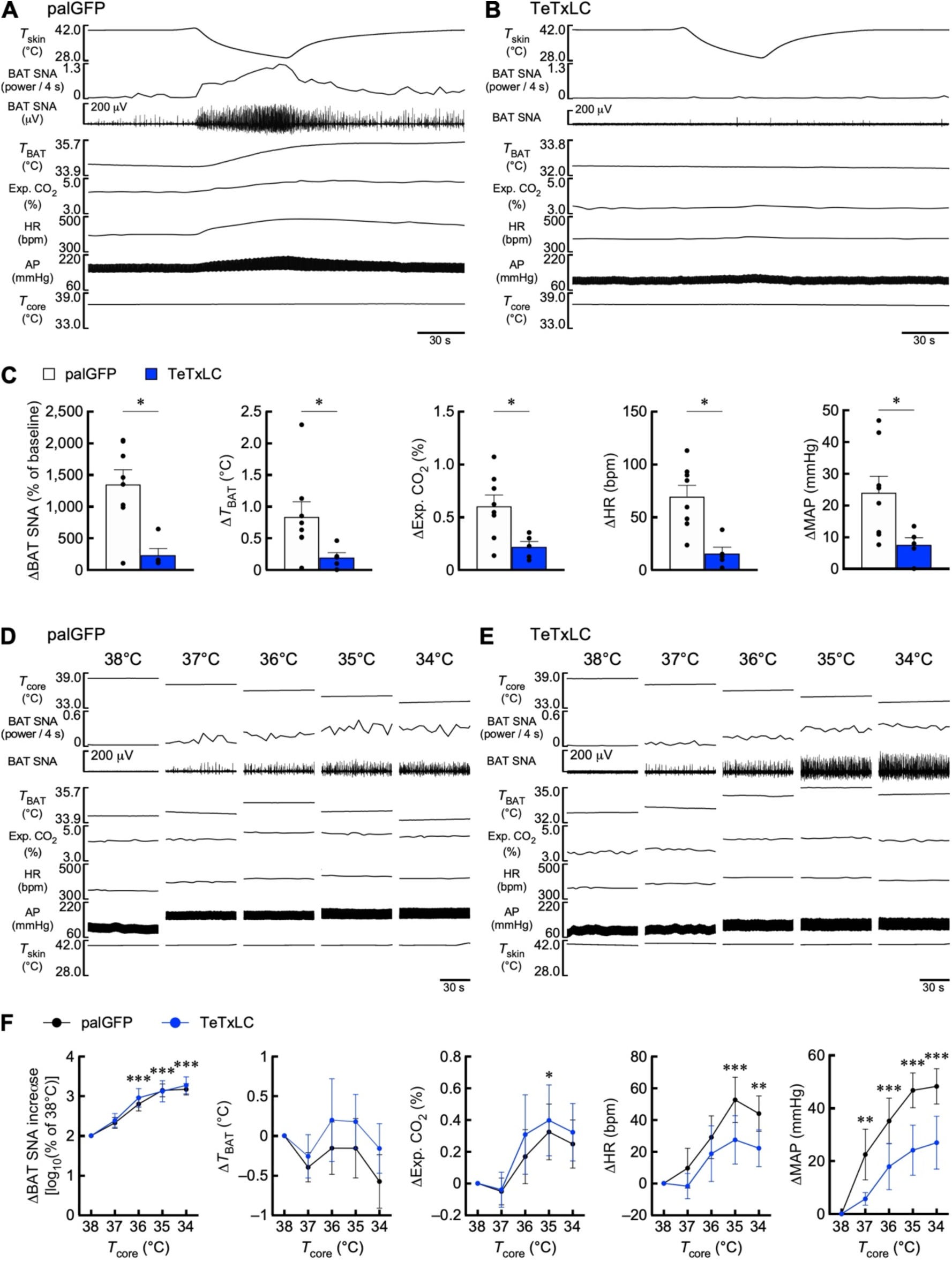
TeTxLC-mediated chronic suppression of LPB→CeA neurons abolishes BAT thermogenic and cardiovascular responses to skin cooling, but not responses to body core cooling. **A and B**, Effects of palGFP- and TeTxLC-expression in LPB→CeA neurons on BAT thermogenic and cardiovascular responses to skin cooling. **C**, Skin cooling-evoked changes in physiological variables, which were analyzed by Mann-Whitney U tests (palGFP: *n* = 8, TeTxLC: *n* = 5; ΔBAT SNA: *U* = 5; Δ*T*_BAT_: *U* = 4; ΔExp. CO_2_: *U* = 4; ΔHR: *U* = 1; ΔMAP: *U* = 5). **D and E**, Effects of palGFP- and TeTxLC-expression in LPB→CeA neurons on the BAT thermogenic and cardiovascular responses to reduction of *T*_core_. **F**, Body core cooling-evoked changes in BAT SNA, *T*_BAT_, Exp. CO_2_, MAP, and HR were compared to the values at 38°C. Data from control (palGFP) rats (*n* = 8) were analyzed by repeated measures one-way ANOVA followed by Bonferroni’s post hoc test (ΔBAT SNA: *F*_4, 28_ = 31.41, *P* < 0.001; Δ*T*_BAT_: *F*_4, 28_ = 1.72, *P* = 0.175; ΔExp. CO_2_: *F*_4, 28_ = 3.67, *P* = 0.016; ΔHR: *F*_4, 28_ = 7.16, *P* < 0.001; ΔMAP: *F*_4, 28_ = 18.25, *P* < 0.001). \**P* < 0.05; \*\**P* < 0.01; \*\*\**P* < 0.001 (vs 38°C). Body cooling-evoked changes in the physiological variables in the TeTxLc group were not significantly different from those in the palGFP group, as determined by Bonferroni’s post hoc test following repeated measures two-way ANOVA (palGFP, *n* = 8; TeTxLC, *n* = 5; ΔBAT SNA: group: *F*_1, 11_ = 0.09, *P* = 0.770, *T*_core_: *F*_4, 44_ = 49.85, *P* < 0.001, interaction: *F*_4, 44_ = 0.25, *P* = 0.911; Δ*T*_BAT_: group: *F*_1, 11_ = 0.44, *P* = 0.519, *T*_core_: *F*_4, 44_ = 2.12, *P* = 0.095, interaction: *F*_4, 44_ = 0.41, *P* = 0.800; ΔExp. CO_2_: group: *F*_1, 11_ = 0.10, *P* = 0.754, *T*_core_: *F*_4, 44_ = 7.20, *P* < 0.001, interaction: *F*_4, 44_ = 0.18, *P* = 0.950; ΔHR: group: *F*_1, 11_ = 1.01, *P* = 0.336, *T*_core_: *F*_4, 44_ = 8.24, *P* < 0.001, interaction: *F*_4, 44_ = 0.67, *P* = 0.619; ΔMAP: group: *F*_1, 11_ = 3.21, *P* = 0.101, *T*_core_: *F*_4, 44_ = 19.68, *P* < 0.001, interaction: *F*_4, 44_ = 1.65, *P* = 0.180). Error bars indicate SEM.

To examine the effects of skin cooling on physiological parameters (Fig. 2A–C and 7A–C), ice-cold water was perfused through the water jacket to lower *T*_skin_ from 40°C to 30°C. For data analyses in Fig. 2C and 7C, baseline values of BAT SNA, *T*_BAT_, expired CO_2_, HR, and mean arterial pressure (MAP) were the averages during the 30-s period immediately before the initiation of skin cooling. Skin cooling-evoked changes in *T*_BAT_, expired CO_2_, HR, and MAP were the differences between the baseline values and their peak values within 30 s (60 s for *T*_BAT_) prior to the end of skin cooling. Skin cooling-evoked changes in BAT SNA were the average of the power/4-s value for the 30-s period before the end of skin cooling and expressed as % of the baseline values.

To examine the effects of *T*_core_ on physiological parameters (Fig. 2D–F and 7D–F), *T*_core_ was gradually reduced from 38°C to 34°C by repeated skin cooling, and physiological parameters were measured for 60 sec each time *T*_core_ was reduced by 1°C and *T*_skin_ was at 40°C. In Fig. 2F and 7F, 60-s average values of BAT SNA, *T*_BAT_, expired CO_2_, HR, and MAP at *T*_core_ of 38°C were taken as baseline values, and differences from the baseline values in 60-s average values of *T*_BAT_, expired CO_2_, HR, and MAP at each *T*_core_ were calculated. A 60-s average of the power/4-s value of BAT SNA at each *T*_core_ was expressed as log_10_% of the baseline values at *T*_core_ of 38°C [log_10_ (% of 38°C BAT SNA)].

After all the recordings, the rats were transcardially perfused and the brain tissue was processed to confirm AAV injections and gene expression as described below.

### Daily monitoring of *T*_core_ and appetitive behaviors and thermal exposure

Diurnal changes in *T*_core_, activity, food intake, and water consumption of rats were measured in a cage equipped with an automated feeding behavior measuring device (Feedam, Melquest, Toyama, Japan) placed in a climate chamber air-conditioned at 25 ± 0.5°C with a 12-h light/dark cycle (light: 7:00–19:00, dark: 19:00–7:00). *T*_core_ and activity were measured by receiving signals from the implanted telemetry transmitter with receiver boards placed near the cage. Rats were individually housed in the cages from the day before the measurement to acclimate them to the environment, and measurements were performed over the next two days. Amounts of food and water consumed were measured every 10 min and used to calculate daily and hourly food intake and water consumption per body weight. To measure *T*_core_, activity, food intake, and water consumption in a cold or hot environment, the air temperature in the climate chamber was set at 4°C or 36°C, respectively, for 2 h (approximately between 9:00 and 15:00).

### Immunohistochemistry

Immunohistochemical procedures followed our previous methods (Nakamura et al., 2000, 2004). Rats were deeply anesthetized and transcardially perfused with 4% formaldehyde in 0.1 M phosphate buffer (pH 7.4). The brain was removed, postfixed in the fixative at 4°C for 2–3 h, and then cryoprotected in a 30% sucrose solution overnight. The tissue was cut into 30-μm-thick frontal sections on a freezing microtome.

For anterograde tracing of palGFP-labeled axons, sections were incubated overnight with an anti-GFP mouse antibody (1:200; A11120, Thermo Fisher Scientific) and then with an Alexa488-conjugated goat antibody to mouse IgG (10 μg/ml; A11029, Thermo Fisher Scientific) for 1 h.

For double immunofluorescence labeling of CTb and Fos, sections were incubated overnight with an anti-CTb goat serum (1:4,000; #703, List Biological Labs) and an anti-Fos rabbit serum (1:10,000; PC38, Merck Millipore). After rinsing, the sections were incubated with an Alexa594-conjugated donkey antibody to goat IgG (5 μg/ml; A11058, Thermo Fisher Scientific) for 1 h and then blocked with 10% normal goat serum for 30 min. After a thorough wash, the sections were incubated with a biotinylated donkey antibody to rabbit IgG (10 μg/ml; AP182B, Merck Millipore) and then with Alexa488-conjugated streptavidin (2.5 μg/ml; S11223, Thermo Fisher Scientific).

For detection of EYFP and Cre recombinase co-expressed in cells AAV-transduced with TeTxLC, sections were incubated overnight with an anti-GFP rabbit antibody (0.5 μg/ml) (Tamamaki et al., 2000) and an anti-Cre recombinase mouse monoclonal antibody (1:1,000; MAB3120, Merck Millipore). After a rinse, the sections were incubated for 1 h with an Alexa488-conjugated goat antibody to rabbit IgG (5 μg/ml; A11034, Thermo Fisher Scientific) and a biotinylated donkey antibody to mouse IgG (10 μg/ml; AP192B, Merck Millipore). The sections were further incubated with Alexa594-conjugated streptavidin (2.5 μg/ml; S11227, Thermo Fisher).

For detection of mCherry and Cre recombinase co-expressed in cells AAV-transduced with hM4Di^nrxn^ or iChloC-mCherry, sections were incubated overnight with an anti-RFP rabbit antibody (0.5 μg/ml; R10367, Thermo Fisher Scientific) and an anti-Cre recombinase mouse monoclonal antibody (1:1,000). After a rinse, the sections were incubated for 1 h with an Alexa594-conjugated goat antibody to rabbit IgG (5 μg/ml; A11012, Thermo Fisher Scientific) and a biotinylated donkey antibody to mouse IgG (10 μg/ml). The sections were further incubated with Alexa488-conjugated streptavidin (2.5 μg/ml). For detection of axon terminals of neurons transduced with iChloC-mCherry, sections were incubated overnight with an anti-monomeric RFP guinea pig antibody (0.1 μg/ml) (Hioki et al., 2010). After a rinse, the sections were incubated for 1 h with a biotinylated donkey antibody to guinea pig IgG (10 μg/ml; 706-065-148, Jackson ImmunoResearch). The sections were further incubated with Alexa594-conjugated streptavidin (2.5 μg/ml).

Stained sections were mounted on glass slides and coverslipped with 50% glycerol/50% PBS containing 2.5% triethylenediamine. The sections were observed under an epifluorescence microscope (Eclipse 80i, Nikon) or a confocal laser scanning microscope (TCS SP8, Leica).

### Anatomy and statistics

The anatomical nomenclature of most brain regions followed the brain atlas of Paxinos and Watson (2007).

Data are presented as the means ± S.E.M. Statistic comparison analyses were performed using a paired or unpaired t-test, a Mann-Whitney U test, a repeated measures one-way ANOVA followed by Bonferroni’s multiple comparisons test, or a repeated measures two-way ANOVA followed by Bonferroni’s multiple comparisons test (Prism 9, GraphPad) as stated in the text and figure legends. All the statistical tests were two-sided. Statistical results with a *P* value of < 0.05 were considered significant.

## RESULTS

### LPB→MnPO neurons are required for heat avoidance but not cold avoidance

To identify candidate forebrain areas to which LPB neurons transmit thermosensory signals, we performed anterograde neural tract tracing from the LPB in rats by unilaterally injecting AAV-CMV-palGFP into the LPB. Expression of palGFP in LPB neurons visualized their axon fibers innervating many, but specific forebrain regions, and particularly dense innervation was observed in the MnPO and CeA (Fig. 1, A and 1B), consistent with earlier observations (Saper and Loewy, 1980; Fulwiler and Saper, 1984). We have previously shown that two segregated groups of neurons in the external lateral part (LPBel) and the dorsal part of the LPB (LPBd) are activated by cutaneous cold and warm sensory inputs and transmit the thermosensory signals to the POA to elicit cold-defensive and heat-defensive autonomous responses, respectively (Nakamura and Morrison, 2008, 2010). Therefore, in the present study, we first investigated whether LPB→MnPO projection neurons are involved in behavioral thermoregulation.

To this end, we chronically inhibited transmission of LPB→MnPO neurons in rats by pathway-selective transduction with TeTxLC (Fig. 1C), which blocks neurotransmitter release at axon terminals by cleaving v-SNARE proteins (Link et al., 1992; Schiavo et al., 1992). An AAV for Cre recombinase-dependent expression of TeTxLC and an AAVrg for Cre expression were injected into the LPB (bilateral) and MnPO, respectively, to selectively inhibit double-infected neurons (*i.e.*, LPB→MnPO neurons) (Fig. 1C). As expected, numerous neurons in the LPBel and LPBd expressed both Cre recombinase and EYFP (indicative of TeTxLC expression) and there were few EYFP-expressing neurons without Cre expression (Fig. 1D). LPB→MnPO neurons in control rats were transduced with palGFP instead of TeTxLC. palGFP-labeled axons of LPB→MnPO neurons were densely distributed in the MnPO and adjacent medial preoptic area, but were not observed in the CeA (Fig. 1E).

The rats were subjected to a two-floor innocuous TPPT (Fig. 1F). During the test, the rats freely moved between the two thermal plate floors, one of which was set at 28°C, the thermoneutral temperature for laboratory rats, and the other was set at 39°C (warm) or 15°C (cold) under a room temperature of 25°C. Control rats preferred to stay on the 28°C plate rather than the 39°C or 15°C plate, exhibiting typical heat and cold avoidance behavior (Fig. 1, G and H). In contrast, TeTxLC-transduced rats failed to exhibit heat avoidance, resulting in a higher *T*_core_ than control rats (Fig. 1G). However, TeTxLC-transduced rats did exhibit cold avoidance like control rats and changes in *T*_core_ were not different between the groups (Fig. 1H). In the tests with both temperature settings, the number of transition times and the distances the rats moved between the plates did not differ between control and TeTxLC-transduced rats (Fig. 1, G and H). These results indicate that heat avoidance, but not cold avoidance, requires transmission by LPB→MnPO neurons.

Because we have shown that cold sensory signals through the LPB→MnPO pathway elicit autonomous cold defense responses (Nakamura and Morrison, 2008), the effect of TeTxLC-mediated blockade of LPB→MnPO neurons on skin cooling-evoked BAT thermogenesis was examined in an anesthetized preparation. In palGFP-transduced control rats, cooling the trunk skin evoked increases in BAT SNA, *T*_BAT_, expired CO_2_, and HR under the conditions of 37°C *T*_core_ (Fig. 2A) (Nakamura and Morrison, 2007). However, these skin cooling-evoked physiological responses were abolished in TeTxLC-transduced rats under the same *T*_core_ conditions (Fig. 2, B and C). Skin cooling-evoked changes in mean arterial pressure (MAP) were comparable between the rat groups (Fig. 2C). These results are consistent with the notion that LPB→MnPO thermosensory transmission is required to elicit feedforward thermogenic and cardiac responses to environmental cold challenges (Nakamura and Morrison, 2008).

Next, we examined the effect of TeTxLC-mediated blockade of LPB→MnPO neurons on BAT thermogenesis evoked by body core cooling. In control rats, reduction of *T*_core_ in 1°C decrements from 38°C to 34°C, while *T*_skin_ was controlled at 40°C, gradually increased BAT SNA, expired CO_2_ and MAP, but *T*_BAT_ and HR were sustained (Fig. 2, D and F). In TeTxLC-transduced rats, however, the same reduction in *T*_core_ did not evoke BAT SNA and significantly reduced *T*_BAT_, expired CO_2_, and MAP compared to control rats (Fig. 2, E and F). These results indicate that LPB→MnPO cold sensory transmission is required not only for physiological responses to environmental cooling, but also for cold defense responses to body cooling. The results also indicate that the AAV-transduction with TeTxLC successfully blocked the cold sensory transmission by LPB→MnPO neurons, supporting our result from the TPPT that these neurons are not involved in cold avoidance.

We further investigated the effect of chronic blockade of LPB→MnPO transmission on the diurnal control of body temperature and appetitive behaviors. TeTxLC-transduced rats maintained a higher *T*_core_ during the dark phase than control rats (Fig. 3A). Daily food intake of TeTxLC-transduced rats was also higher than that of control rats, although there was no obvious difference in circadian changes in hourly food intake (Fig. 3C). The level of activity and water intake were comparable between the rat groups (Fig. 3, B and D). To test the ability of TeTxLC-transduced rats to defend *T*_core_ against ambient heat and cold, they were exposed to an environment of 36°C or 4°C for 120 min. During exposure to 36°C, *T*_core_ of TeTxLC-transduced rats showed a greater increase than that of control rats, reaching almost 40°C at the end of the exposure (Fig. 3E). Activity counts, food intake, and water intake during heat exposure were comparable between the rat groups (Fig. 3, F to H). During exposure to 4°C, TeTxLC-transduced rats exhibited a significant reduction in *T*_core_ compared to control rats, which were able to maintain their *T*_core_ around 37°C (Fig. 3I). Control rats exhibited a cold-induced increase in food intake, which supplies energy for adaptive thermogenesis, during the second half of the cold exposure, whereas TeTxLC-transduced rats showed a reduced food intake (Fig. 3K). Activity and water intake during the cold exposure did not differ between the rat groups (Fig. 3, J and L). These results demonstrate that thermosensory afferent signals transmitted by LPB→MnPO neurons are essential for maintaining thermal homeostasis in hot and cold environments.

**Figure 3.**
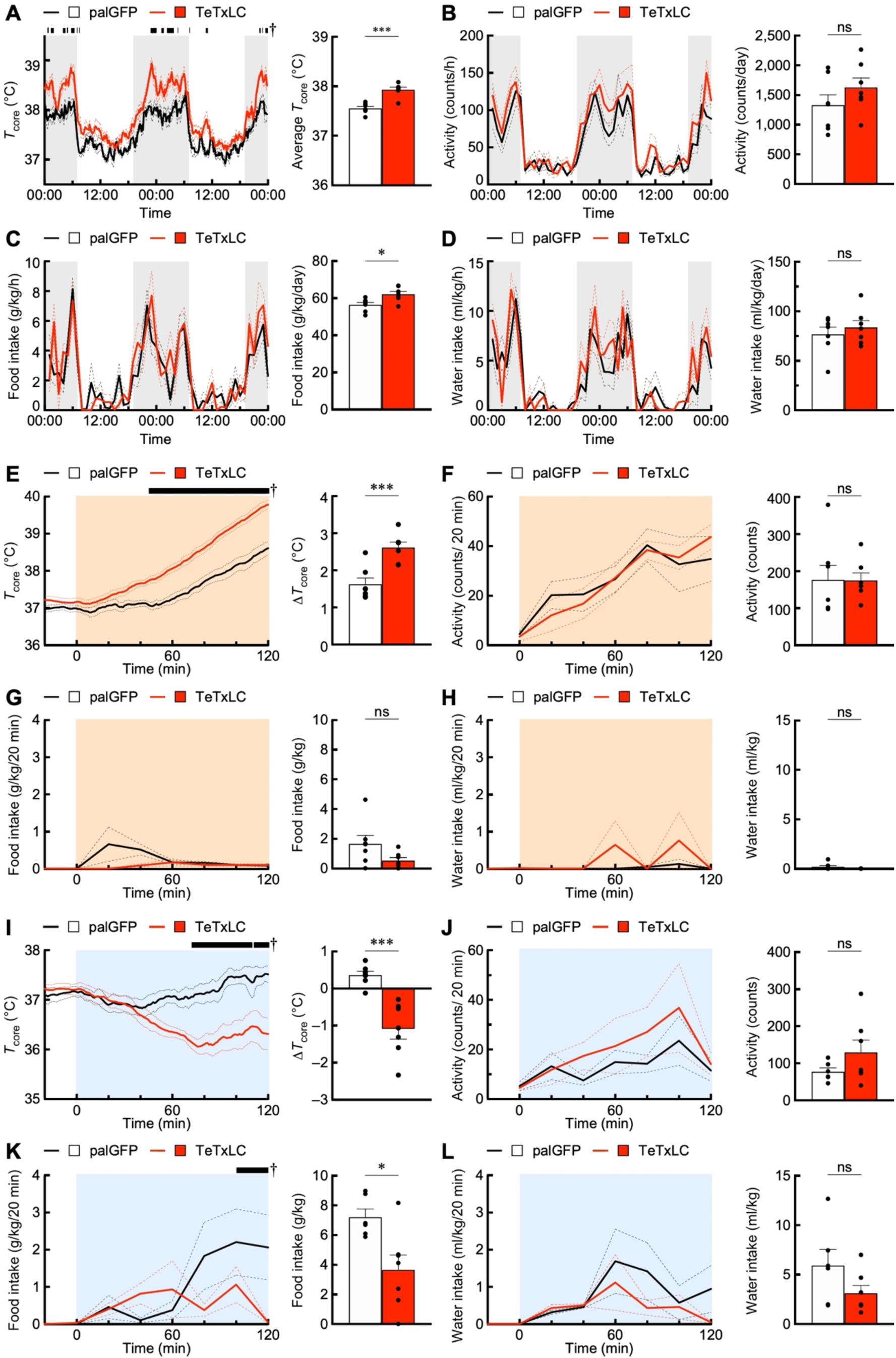
TeTxLC-mediated suppression of LPB→MnPO neurons affects diurnal control of body temperature and appetitive behaviors and heat and cold tolerance. **A–D**, Effects of TeTxLC-expression in LPB→MnPO neurons on diurnal changes in *T*_core_, activity counts, food intake, and water intake (left), which were analyzed by repeated measures two-way ANOVA followed by Bonferroni’s post hoc test [*n* = 7 per group; (A): group: *F*_1, 12_ = 35.17, *P* < 0.001, time: *F*_287, 3444_ = 38.10, *P* < 0.001, interaction: *F*_287, 3444_ = 2.03, *P* < 0.001; (B): group: *F*_1, 12_ = 1.65, *P* = 0.223, time: *F*_47, 564_ = 20.87, *P* < 0.001, interaction: *F*_47, 564_ = 0.84, *P* = 0.771; (C): group: *F*_1, 12_ = 8.42, *P* = 0.013, time: *F*_47, 564_ = 12.45, *P* < 0.001, interaction: *F*_47, 564_ = 1.24, *P* = 0.139; (D): group: *F*_1, 12_ = 0.52, *P* = 0.487, time: *F*_47, 564_ = 13.75, *P* < 0.001, interaction: *F*_47, 564_ = 1.47, *P* = 0.026]. Time points with a significant difference between the groups (*P* < 0.05) are indicated by black bars with †. Right graphs show intergroup differences in average *T*_core_ and daily activity counts, food intake and water intake were analyzed by unpaired *t*-tests (*n* = 7 per group; (A) *t*_12_ = 5.92; (B) *t*_12_ = 1.29; (C) *t*_12_ = 2.90; (D) *t*_12_ = 0.72). \**P* < 0.05; \*\*\**P* < 0.001; ns, not significant. **E–L**, Rats with palGFP- or TeTxLC-expression in LPB→MnPO neurons were exposed to 36°C (E–H) or 15°C (I–L). Time-course changes in *T*_core_, activity counts, food intake, and water intake (left graphs) were analyzed by repeated measures two-way ANOVA followed by Bonferroni’s post hoc test [*n* = 7 per group; (E): group: *F*_1, 12_ = 23.98, *P* < 0.001, time: *F*_140, 1680_ = 141.0, *P* < 0.001, interaction: *F*_140, 1680_ = 10.69, *P* < 0.001; (F): group: *F*_1, 12_ = 0.00, *P* = 0.953, time: *F*_6, 72_ = 17.52, *P* < 0.001, interaction: *F*_6, 72_ = 0.71, *P* = 0.643; (G): group: *F*_1, 12_ = 3.49, *P* = 0.087, time: *F*_7, 84_ = 2.03, *P* = 0.061, interaction: *F*_7, 84_ = 2.21, *P* = 0.041; (H): group: *F*_1, 12_ = 0.78, *P* = 0.393, time: *F*_7, 84_ = 1.17, *P* = 0.331, interaction: *F*_7, 84_ = 0.81, *P* = 0.579; (I): group: *F*_1, 12_ = 7.84, *P* = 0.016, time: *F*_140, 1680_ = 4.296, *P* < 0.001, interaction: *F*_140, 1680_ = 7.68, *P* < 0.001; (J): group: *F*_1, 12_ = 1.473, *P* = 0.248, time: *F*_6, 72_ = 2.42, *P* = 0.034, interaction: *F*_6, 72_ = 0.36, *P* = 0.904; (K): group: *F*_1, 12_ = 9.27, *P* = 0.010, time: *F*_7, 84_ = 2.69, *P* = 0.014, interaction: *F*_7, 84_ = 2.08, *P* = 0.055; (L): group: *F*_1, 12_ = 2.11, *P* = 0.171, time: *F*_7, 84_ = 2.60, *P* = 0.018, interaction: *F*_7, 84_ = 0.61, *P* = 0.746]. Time points with a significant difference between the groups (*P* < 0.05) are indicated by black bars with †. Right graphs show intergroup differences in Δ*T*_core_ (difference between values at time 0 and 120 min), total activity counts, total food intake, and water intake, which were analyzed by unpaired *t*-tests [*n* = 7 per group; (E) *t*_12_ = 4.51; (F) *t*_12_ = 0.03; (G) *t*_12_ = 1.90; (H) *t*_12_ = 1.39; (I) *t*_12_ = 4.96; (J) *t*_12_ = 1.23; (K) *t*_12_ = 3.01; (L) *t*_12_ = 1.38]. \**P* < 0.05; \*\*\**P* < 0.001; ns, not significant. Error bars indicate SEM.

### LPB→MnPO neurons mediate heat avoidance via synaptic inputs to the MnPO

It was possible that the TeTxLC-mediated chronic inhibition of LPB→MnPO neurons might have abolished heat avoidance by causing long-term circuit alternations. Moreover, axons of LPB→MnPO neurons were distributed not only in the POA including the MnPO, but their collaterals were densely distributed in several other regions including the DMH, paraventricular hypothalamic nucleus, lateral hypothalamic area (LH), paraventricular thalamic nucleus, parasubthalamic nucleus, and lateral periaqueductal gray (Fig. 1E). Therefore, we performed acute local chemogenetic inhibition of synaptic release to examine whether direct synaptic inputs from the LPB to the MnPO mediate the thermosensory afferent signaling required to elicit heat avoidance. The Cre-dependent expression system with anterograde and retrograde AAVs was used to selectively transduce LPB→MnPO neurons with the inhibitory DREADD hM4Di^nrxn^, an axon-targeted variant of hM4Di conjugated with an intracellular domain of neurexin 1α (Stachniak et al., 2014) (Fig. 4A). Activation of hM4Di^nrxn^ suppresses synaptic release probability (Stachniak et al., 2014). Similar to TeTxLC expression (Fig. 1C), many neurons in the LPBel and LPBd expressed both Cre and mCherry (indicative of hM4Di^nrxn^ expression) and there were few mCherry-expressing neurons without Cre expression (Fig. 4B). LPB→MnPO neurons in control rats were transduced with palGFP instead of hM4Di^nrxn^.

**Figure 4.**
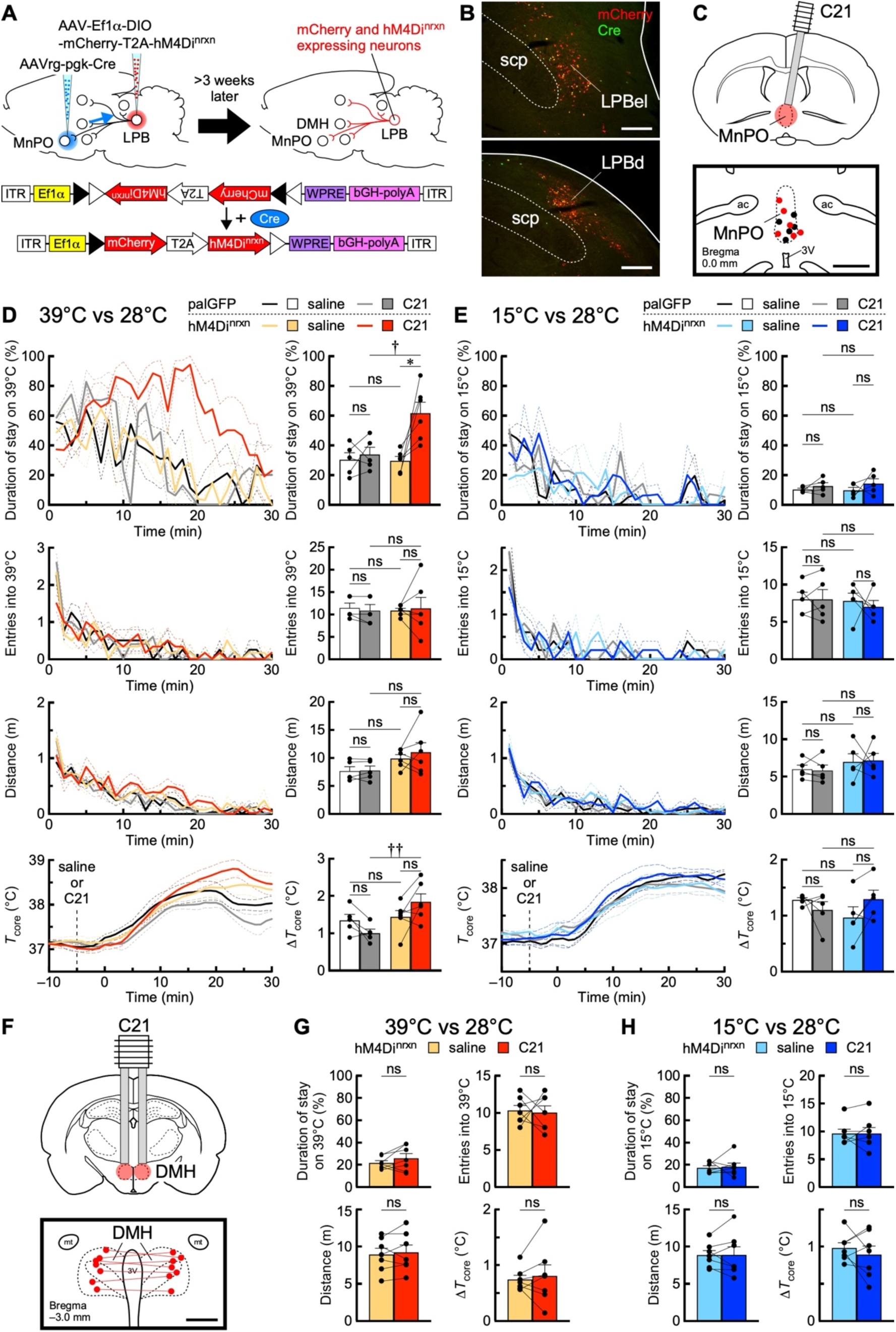
Local chemogenetic inhibition of presynaptic release from MnPO axon terminals of LPB→MnPO neurons abolishes heat avoidance. **A**, Selective transduction of LPB→MnPO neurons with hM4Di^nrxn^-T2A-mCherry using AAV and AAVrg. Control rats expressed palGFP instead of hM4Di^nrxn^-T2A-mCherry. **B**, Cre- and/or mCherry-immunoreactive neurons in the LPB. Scale bars, 500 μm. **C**, The AAV-injected rats were microinjected with saline or C21 into the MnPO (top). Bottom map shows injection sites in the MnPO of the rats AAV-transduced with palGFP (black dots) or mCherry-T2A-hM4Di^nrxn^ (red dots). Scale bar, 1.0 mm. **D and E**, Effects of saline or C21 microinjection into the MnPO of rats expressing palGFP or hM4Di^nrxn^ in LPB→MnPO neurons on heat (D) and cold (E) avoidance. Time-course changes (left, every 1 min) in % duration of stay on 39°C or 15°C plate, times of entry into 39°C or 15°C plate, distance traveled, and *T*_core_. Right graphs show % duration of stay on 39°C or 15°C plate, total times of entry into 39°C or 15°C plate, total distance traveled, and Δ*T*_core_ (difference between value at time 0 and peak) for the 30-min test period, and the data were analyzed by repeated measures one-way ANOVA followed by Bonferroni’s post hoc test [(D): palGFP, *n* = 5; hM4Di^nrxn^, *n* = 6; duration of stay: *F*_3, 18_ = 8.87, *P* = 0.001; entries: *F*_3, 18_ = 0.04, *P* = 0.989; distance: *F*_3, 18_ = 2.25, *P* = 0.117; Δ*T*_core_: *F*_3, 18_ = 4.09, *P* = 0.022; (E): palGFP, *n* = 5; hM4Di^nrxn^, *n* = 5; duration of stay: *F*_3, 16_ = 0.91, *P* = 0.456; entries: *F*_3, 16_ = 0.21, *P* = 0.889; distance: *F*_3, 16_ = 0.66, *P* = 0.588; Δ*T*_core_: *F*_3, 16_ = 1.17, *P* = 0.353]. \**P* < 0.05; ^†^*P* < 0.05; ^††^*P* < 0.01; ns, not significant. **F**, Rats expressing mCherry-T2A-hM4Di^nrxn^ in LPB→MnPO neurons received bilateral microinjections into the DMH with saline or C21 (top). Bottom map shows injection sites (red dots tied with a line). Scale bar, 500 μm. mt, mammillothalamic tract. **G and H**, Effects of saline or C21 microinjections into the DMH on heat (G) and cold (H) avoidance. Graphs show % duration of stay on 39°C or 15°C plate, total times of entry into 39°C or 15°C plate, total distance traveled, and Δ*T*_core_ (difference between value at time 0 and peak) for the 30-min test period, and the data were analyzed by paired *t*-test [*n* = 7; (G): duration of stay, *t*_6_ = 1.30; entries, *t*_6_ = 0.22; distance, *t*_6_ = 0.50; Δ*T*_core_, *t*_6_ = 0.47; (H): duration of stay, *t*_6_ = 0.26; entries, *t*_6_ = 0.00; distance, *t*_6_ = 0.05; Δ*T*_core_, *t*_6_ = 0.83]. ns, not significant. Error bars indicate SEM.

To locally inhibit synaptic release from LPB→MnPO axons in the MnPO, we nanoinjected the actuator, DREADD agonist 21 (C21), through a cannula preimplanted into the MnPO 5 min before TPPT (Fig. 4C). When tested for heat avoidance (39°C vs 28°C), hM4Di^nrxn^-transduced rats nanoinjected with C21 into the MnPO failed to exhibit heat avoidance, whereas the same rats did exhibit heat avoidance following saline nanoinjection (Fig. 4D). In contrast, control rats nanoinjected with either saline or C21 into the MnPO exhibited heat avoidance. Due to the lack of heat avoidance, hM4Di^nrxn^-transduced rats exhibited a significantly greater increase in *T*_core_ following C21 nanoinjection than control rats, which also exhibited a mild elevation of *T*_core_ due to injection stress (Fig. 4D). C21 did not affect the number of transition times or the distances the rats moved between the plates (Fig. 4D). In the cold avoidance test (15°C vs 28°C), nanoinjection of C21 did not affect cold avoidance or changes in *T*_core_ exhibited by hM4Di^nrxn^-transduced rats (Fig. 4E). In another group of rats, we inhibited synaptic release in the DMH from collateral axons of LPB→MnPO neurons and examined the effect on heat and cold avoidance. Bilateral nanoinjections of C21 into the DMH of rats with hM4Di^nrxn^-expression in LPB→MnPO neurons did not affect heat or cold avoidance behavior or *T*_core_ changes (Fig. 4, F–H). This result means that C21 injected locally at the DMH did not diffuse to the MnPO to affect the LPB→MnPO axon terminals, indicating a localized effect of C21. These results demonstrate that thermosensory synaptic inputs to the MnPO from LPB→MnPO neurons mediate heat avoidance.

### LPB→CeA neurons are cold-activated and segregate from LPB→MnPO neurons

To investigate the involvement of LPB→CeA neurons in innocuous thermosensory transmission, we performed a functional retrograde tracing study. CTb, a retrograde neural tracer, was bilaterally injected into the CeA (Fig. 5, A and B), and the rats were exposed to an environment of 4°C (cold exposure), 25°C (control exposure) or 36°C (heat exposure) for 2 h. Among the rat groups, comparable numbers of CTb-labeled cell bodies were observed in the parabrachial nucleus subdivisions: the central part of the LPB, the LPBd, the LPBel, the internal part of the LPB, and the medial parabrachial nucleus (Fig. 5, C–E). Cold exposure significantly increased CTb-labeled neurons with expression of Fos, a marker for neuronal activation, in the LPBel, compared to control exposure (Fig 5, C, D and F). However, heat exposure did not increase Fos expression in CTb-labeled parabrachial neurons (Fig 5, C, D and F).

**Figure 5.**
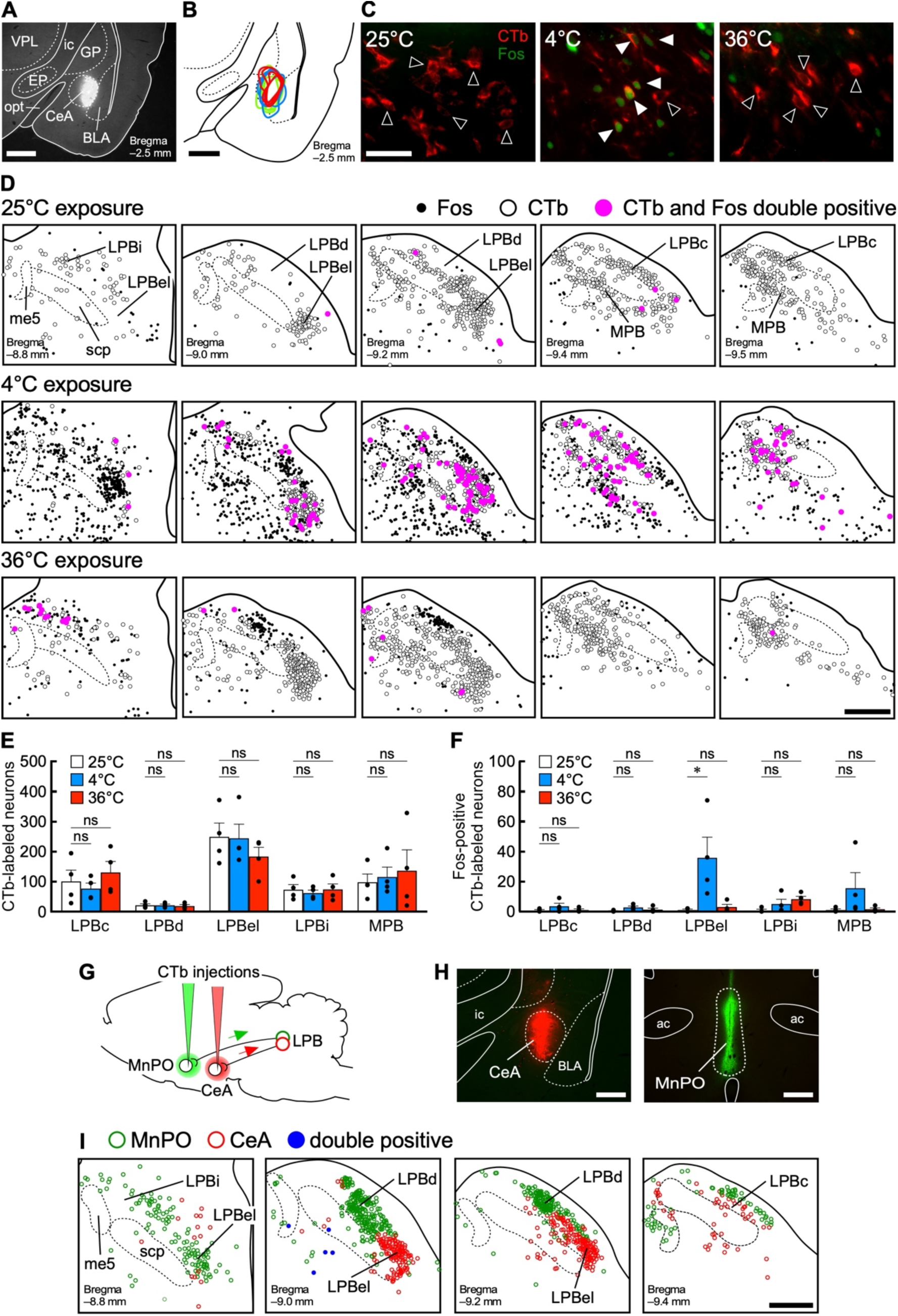
LPB→CeA neurons are cold-activated and segregate from LPB→MnPO neurons. **A and B**, CTb was injected into the CeA. Injection sites in the right CeA for the group data in (E) and (F) are mapped in (B) (green: 25°C-exposed; blue: 4°C-exposed; red: 36°C-exposed). **C**, CTb- and Fos-immunoreactive cells in the LPBel following a 2-h exposure to ambient temperature of 25°C (control), 4°C (cold), or 36°C (heat). Solid and hollow arrowheads indicate CTb-labeled neuronal cell bodies with and without Fos-immunoreactivity, respectively. Scale bar, 50 μm. **D**, Distributions of CTb-labeled and/or Fos-immunoreactive cells in the parabrachial nucleus following exposure to the respective temperature. Scale bar, 500 μm. **E and F**, Numbers of CTb-labeled cells (E) and Fos-immunoreactive CTb-labeled cells (F) in subareas of the parabrachial nucleus (counted in the right hemisphere; *n* = 4 rats per group). Data were analyzed by one-way ANOVA followed by Bonferroni’s post hoc test [(E): LPBc: *F*_2, 9_ = 0.72, *P* = 0.515; LPBd: *F*_2, 9_ = 0.11, *P* = 0.900; LPBel: *F*_2, 9_ = 0.77, *P* = 0.490; LPBi: *F*_2, 9_ = 0.19, *P* = 0.827; MPB: *F*_2, 9_ = 0.17, *P* = 0.849; (F): LPBc: *F*_2, 9_ = 1.55, *P* = 0.264; LPBd: *F*_2, 9_ = 2.82, *P* = 0.112; LPBel: *F*_2, 9_ = 6.00, *P* = 0.022; LPBi: *F*_2, 9_ = 3.19, *P* = 0.090; MPB: *F*_2, 9_ = 1.87, *P* = 0.209]. \**P* < 0.05; ns, not significant. Error bars indicate SEM. **G**, Double retrograde labeling of LPB→MnPO and LPB→CeA neurons with CTb conjugated with different fluorophores. **H**, Injection sites of CTb into the CeA and the MnPO. Scale bars, 500 μm. ac, anterior commissure. **I**, Distribution of CTb-labeled cells in the parabrachial nucleus. Scale bar, 500 μm.

We next compared the distribution of LPB→MnPO neurons and LPB→CeA neurons within the LPB. Injections of CTb with different fluorophores into the MnPO and CeA of the same rat revealed adjacent but clearly segregated distributions of LPB→MnPO neurons and LPB→CeA neurons, and few neurons were double-labeled (Fig. 5, G–I). This result is consistent with the lack of axon collaterals of LPB→MnPO neurons in the CeA (Fig. 1E). In the LPBel, its rostral part harbored LPB→MnPO neurons, which are activated by cutaneous cold sensory inputs (Nakamura and Morrison, 2008), whereas the caudal part was occupied by LPB→CeA neurons, which were also activated by cold exposure (Fig. 5, D and I). These results indicate that distinct groups of LPB neurons innervate the MnPO and CeA, and CeA-projecting neurons in the LPBel are cold-activated.

### Essential roles of LPB→CeA neurons in behavioral and autonomous thermoregulation

To investigate the involvement of LPB→CeA neurons in behavioral thermoregulation, we transduced LPB→CeA neurons with TeTxLC or palGFP (control) by using the Cre-dependent expression system with anterograde and retrograde AAVs (Fig. 6A), and the rats were then subjected to TPPTs. Many neurons expressing both Cre and EYFP (indicative of TeTxLC expression) were distributed in the LPBel (Fig. 6B), similar to neurons labeled with CTb injected into the CeA (Fig. 5D). There were few EYFP-expressing neurons without Cre expression. In TPPTs, control rats preferred the 28°C plate to the 39°C and 15°C plates, indicating avoidance of heat and cold, respectively (Fig. 6, C and D). In contrast, TeTxLC-transduced rats exhibited neither heat nor cold avoidance (Fig. 6, C and D). Due to the failure of heat avoidance, TeTxLC-transduced rats showed a greater increase in *T*_core_ than control rats, but the failure of cold avoidance did not affect *T*_core_ (Fig. 6, C and D). The number of transition times and the distances the rats moved between the plates did not differ between control and TeTxLC-transduced rats (Fig 6, C and D). palGFP-labeled axons of LPB→CeA neurons were abundantly distributed not only in the CeA, but also in several other brain regions including the lateral part of the bed nucleus of the stria terminalis, paraventricular hypothalamic nucleus, VMH, paraventricular thalamic nucleus, parasubthalamic nucleus, and ventrolateral and lateral periaqueductal gray (Fig. 6E). Only sparse axon collaterals of LPB→CeA neurons were observed in the POA, consistent with their segregation from LPB→MnPO neurons. These results suggest that LPB→CeA neurons play an essential role in heat and cold avoidance by transmitting cold sensory signals to these projection sites.

**Figure 6.**
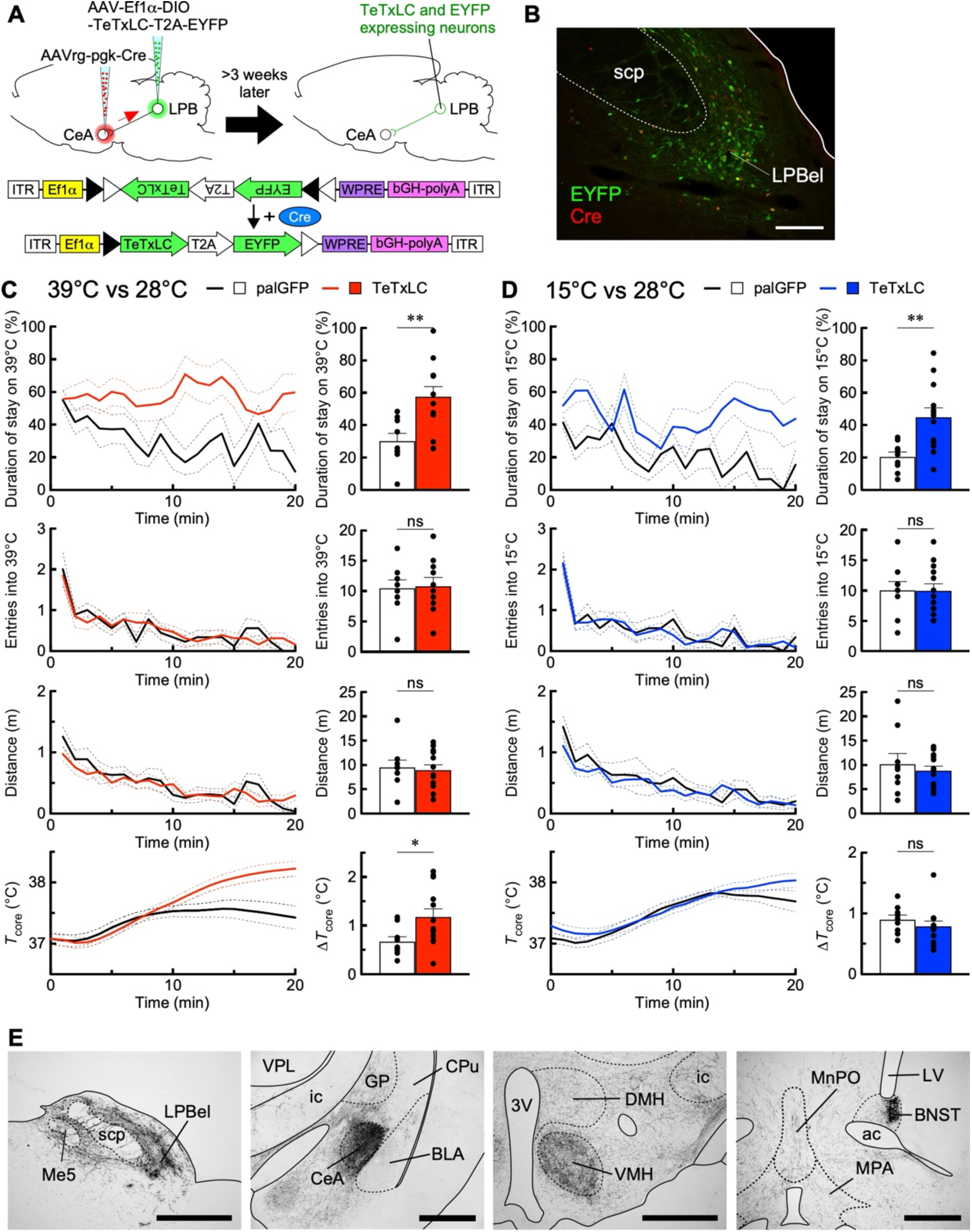
TeTxLC-mediated chronic suppression of LPB→CeA neurons abolishes heat and cold avoidance. **A**, Selective transduction of LPB→CeA neurons with EYFP-T2A-TeTxLC using AAV and AAVrg. **B**, Cre- and/or EYFP-immunoreactive neurons in the LPBel. Scale bar, 500 μm. **C and D**, Effects of the TeTxLC-expression in LPB→CeA neurons on heat (C) and cold (D) avoidance. Control rats expressed palGFP instead of EYFP-T2A-TeTxLC. Time-course changes (left, every 1 min) in % duration of stay on 39°C or 15°C plate, times of entry into 39°C or 15°C plate, distance traveled and *T*_core_. Right graphs show % duration of stay on 39°C or 15°C plate, total times of entry into 39°C or 15°C plate, total distance traveled, and Δ*T*_core_ (difference between value at time 0 and peak) for the 20-min test period, and the data were analyzed by unpaired *t*-tests [palGFP: *n* = 9, TeTxLC: *n* = 13; (C): duration of stay, *t*_20_ = 3.25; entries, *t*_20_ = 0.16; distance, *t*_20_ = 0.32; Δ*T*_core_, *t*_20_ = 2.34; (D): duration of stay, *t*_20_ = 3.32; entries, *t*_20_ = 0.04; distance, *t*_20_ = 0.62; Δ*T*_core_, *t*_20_ = 0.89]. \**P* < 0.05; \*\**P* < 0.01; ns, not significant. Error bars indicate SEM. **E**, Distribution of palGFP-labeled axons of LPB→CeA neurons in a control rat. Scale bars, 1.0 mm. LV, lateral ventricle.

To investigate whether LPB→CeA neurons are also involved in autonomous thermoregulation, we performed *in vivo* electrophysiology in anesthetized rats expressing TeTxLC or palGFP (control) in LPB→CeA neurons. In response to trunk skin cooling under the *T*_core_ condition of 37°C, control rats showed increases in BAT SNA, *T*_BAT_, expired CO_2_, HR, and MAP, which were, however, diminished in TeTxLC-transduced rats (Fig. 7, A–C). In response to reduction of *T*_core_ in 1°C decrements from 38°C to 34°C with *T*_skin_ controlled at 40°C, control rats showed gradual increases in BAT SNA, expired CO_2_, HR, and MAP (Fig. 7, D and F). Interestingly, TeTxLC-transduced rats showed comparable increases in these variables in response to the same reduction in *T*_core_ (Fig. 7, E and F), unlike rats expressing TeTxLC in LPB→MnPO neurons (Fig. 2, E and F). These results indicate that LPB→CeA neurons play an important role not only in behavioral thermoregulation, but also in feedforward autonomous responses to skin cooling.

Further investigation of the effect of chronic blockade of LPB→CeA neurons with TeTxLC on the diurnal control of body temperature and appetitive behaviors revealed that TeTxLC-transduced rats exhibited an approximately 0.5°C higher *T*_core_ than palGFP-transduced control rats throughout the 2-day recording period (Fig. 8A). Daily food intake of TeTxLC-transduced rats was also higher, whereas their activity and water intake were comparable to those of control rats (Fig. 8, B–D). Next, we examined whether the TeTxLC-transduced rats could maintain their *T*_core_ when exposed to a 36°C or 4°C environment for 120 min. During exposure to 36°C, TeTxLC-transduced rats showed a greater increase in *T*_core_ than that of control rats, reaching almost 40°C at the end of exposure (Fig. 8E). Activity counts and food and water intake were not different between the two groups of rats (Fig. 8, F–H). During exposure to 4°C, TeTxLC-transduced rats showed a 1.6°C drop in *T*_core_ and consumed less food and water than control rats, but had comparable activity levels (Fig. 8, I–L). These results show that LPB→CeA neurons play an essential role in the maintenance of body temperature under environmental thermal challenges.

**Figure 8.**
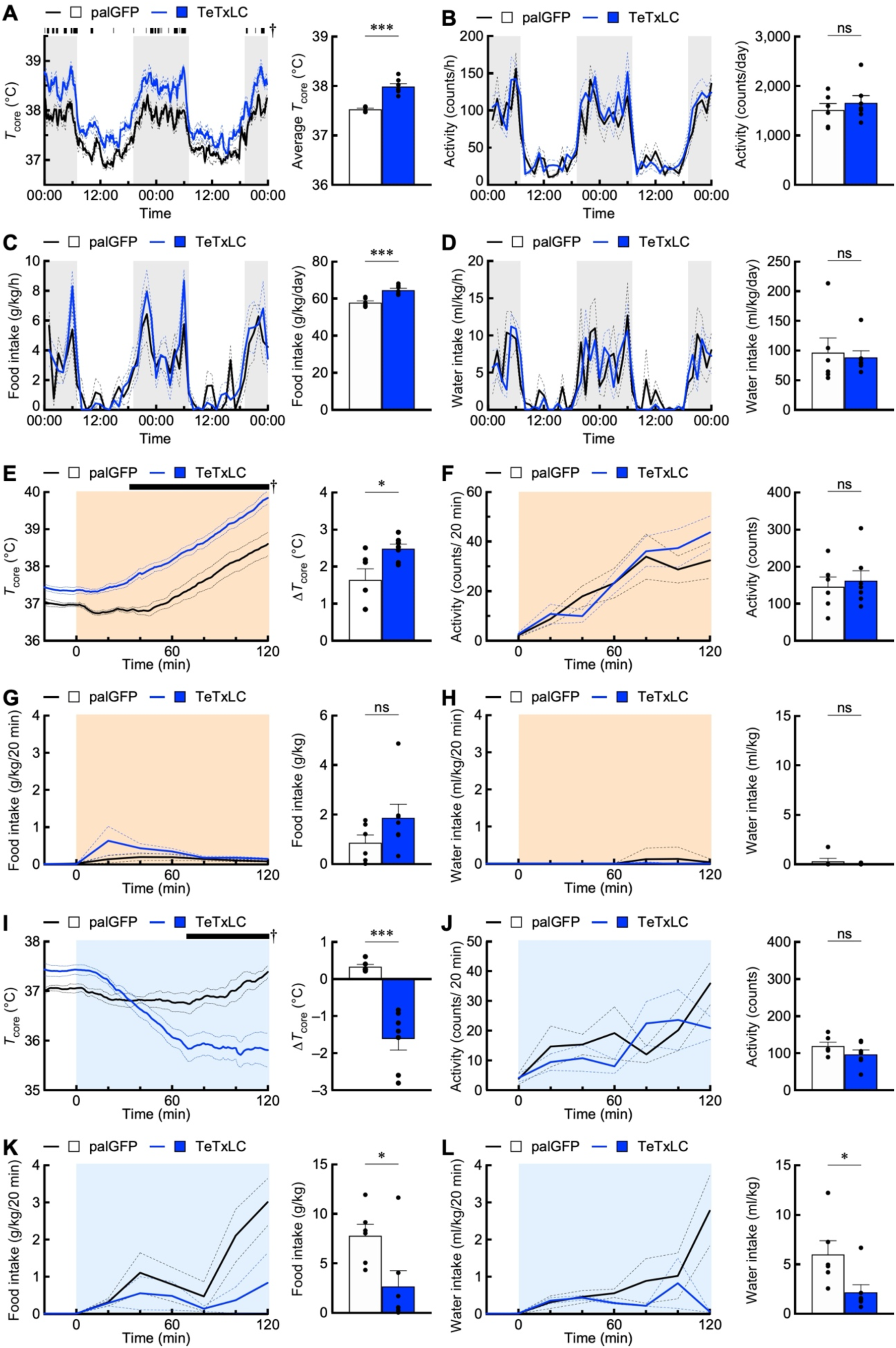
TeTxLC-mediated suppression of LPB→CeA neurons affects diurnal control of body temperature and appetitive behaviors and heat and cold tolerance. **A–D**, Effects of TeTxLC-expression LPB→CeA neurons on diurnal changes in *T*_core_, activity counts, food intake, and water intake (left), which were analyzed by repeated measures two-way ANOVA followed by Bonferroni’s post hoc test [palGFP, *n* = 6; TeTxLC, *n* = 7; (A): group: *F*_1, 11_ = 51.41, *P* < 0.001, time: *F*_287, 3157_ = 35.67, *P* < 0.001, interaction: *F*_287, 3157_ = 1.66, *P* < 0.001; (B): group: *F*_1, 11_ = 0.54, *P* = 0.476, time: *F*_47, 517_ = 20.63, *P* < 0.001, interaction: *F*_47, 517_ = 1.16, *P* = 0.230; (C): group: *F*_1, 11_ = 11.27, *P* = 0.006, time: *F*_47, 517_ = 10.54, *P* < 0.001, interaction: *F*_47, 517_ = 0.83, *P* = 0.784; (D): group: *F*_1, 11_ = 0.08, *P* = 0.786, time: *F*_47, 517_ = 8.41, *P* < 0.001, interaction: *F*_47, 517_ = 1.20, *P* = 0.182]. Time points with a significant difference between the groups (*P* < 0.05) are indicated by black bars with †. Right graphs show intergroup differences in average *T*_core_ and daily activity counts, food intake and water intake were analyzed by unpaired *t*-tests (palGFP, *n* = 6; TeTxLC, *n* = 7; (A) *t*_11_ = 7.114; (B) *t*_11_ = 0.737; (C) *t*_11_ = 5.144; (D) *t*_11_ = 0.317). \*\*\**P* < 0.001; ns, not significant. **E-L**, Rats with palGFP- or TeTxLC-expression in LPB→CeA neurons were exposed to 36°C (E–H) or 15°C (I–L). Time-course changes in *T*_core_, activity counts, food intakes, and water intake (left graphs) were analyzed by repeated measures two-way ANOVA followed by Bonferroni’s post hoc test [palGFP, *n* = 6; TeTxLC, *n* = 7; (E): group: *F*_1, 11_ = 23.73, *P* = 0.005, time: *F*_140, 1540_ = 104.3, *P* < 0.001, interaction: *F*_140, 1540_ = 4.76, *P* < 0.001; (F): group: *F*_1, 11_ = 0.20, *P* = 0.665, time: *F*_6, 66_ = 22.41, *P* < 0.001, interaction: *F*_6, 66_ = 1.14, *P* = 0.348; (G): group: *F*_1, 11_ = 2.46, *P* = 0.145, time: *F*_7, 77_ = 2.98, *P* = 0.008, interaction: *F*_7, 77_ = 1.12, *P* = 0.358; (H): group: *F*_1, 11_ = 1.07, *P* = 0.324, time: *F*_7, 77_ = 1.31, *P* = 0.258, interaction: *F*_7, 77_ = 1.06, *P* = 0.401; (I): group: *F*_1, 11_ = 5.32, *P* = 0.042, time: *F*_140, 1540_ = 15.72, *P* < 0.001, interaction: *F*_140, 1540_ = 15.45, *P* < 0.001; (J): group: *F*_1, 11_ = 2.01, *P* = 0.184, time: *F*_6, 66_ = 3.46, *P* = 0.005, interaction: *F*_6, 66_ = 1.08, *P* = 0.387; (K): group: *F*_1, 11_ = 6.48, *P* = 0.027, time: *F*_7, 77_ = 5.678, *P* < 0.001, interaction: *F*_7, 77_ = 2.30, *P* = 0.035; (L): group: *F*_1, 11_ = 6.32, *P* = 0.029, time: *F*_7, 77_ = 3.61, *P* = 0.002, interaction: *F*_7, 77_ = 3.57, *P* = 0.002]. Time points with a significant difference between the groups (*P* < 0.05) are indicated by black bars with †. Right graphs show intergroup differences in Δ*T*_core_ (difference between values at time 0 and 120 min), total activity counts, food intake, and water intake, which were analyzed by unpaired *t*-tests (palGFP, *n* = 6; TeTxLC, *n* = 7; (E) *t*_11_ = 2.80; (F) *t*_11_ = 0.43; (G) *t*_11_ = 1.54; (H) *t*_11_ = 1.01; (I) *t*_11_ = 5.96; (J) *t*_11_ = 1.41; (K) *t*_11_ = 2.55; (L) *t*_11_ = 2.51). \**P* < 0.05; \*\*\**P* < 0.001; ns, not significant. Error bars indicate SEM.

### LPB→CeA neurons contribute to cold avoidance via axonal inputs to the CeA

Because LPB→CeA neurons project their collateral axons to several brain sites other than the CeA (Fig. 6E), we determined whether their axonal inputs to the CeA mediate heat and cold avoidance behavior by selectively suppressing their CeA axon terminals with an optogenetic technique. The Cre-dependent expression system with anterograde and retrograde AAVs was used to selectively transduce LPB→CeA neurons with iChloC, a chloride-conducting channelrhodopsin for photoinhibition of neuronal activity (Wietek et al., 2015) (Fig. 9A). Both Cre and mCherry-conjugated iChloC were expressed in neurons in the LPBel and there were few mCherry-expressing neurons without Cre expression (Fig. 9B). Even with bilateral local photoinhibition of CeA axon terminals of LPB→CeA neurons over the 20 min period of TPPT (Fig. 9C), iChloC-mCherry-transduced rats exhibited heat avoidance and *T*_core_ changes similar to control rats expressing palGFP instead of iChloC-mCherry (Fig. 9D). In the cold avoidance test, however, local photoinhibition of LPB→CeA axon terminals significantly prolonged the stay of iChloC-mCherry-transduced rats on the 15°C plate, which was particularly evident in the last quarter of the test period, although their *T*_core_ changes and the number of transition times and the distances they moved between the plates were comparable to control rats (Fig. 9E).

**Figure 9.**
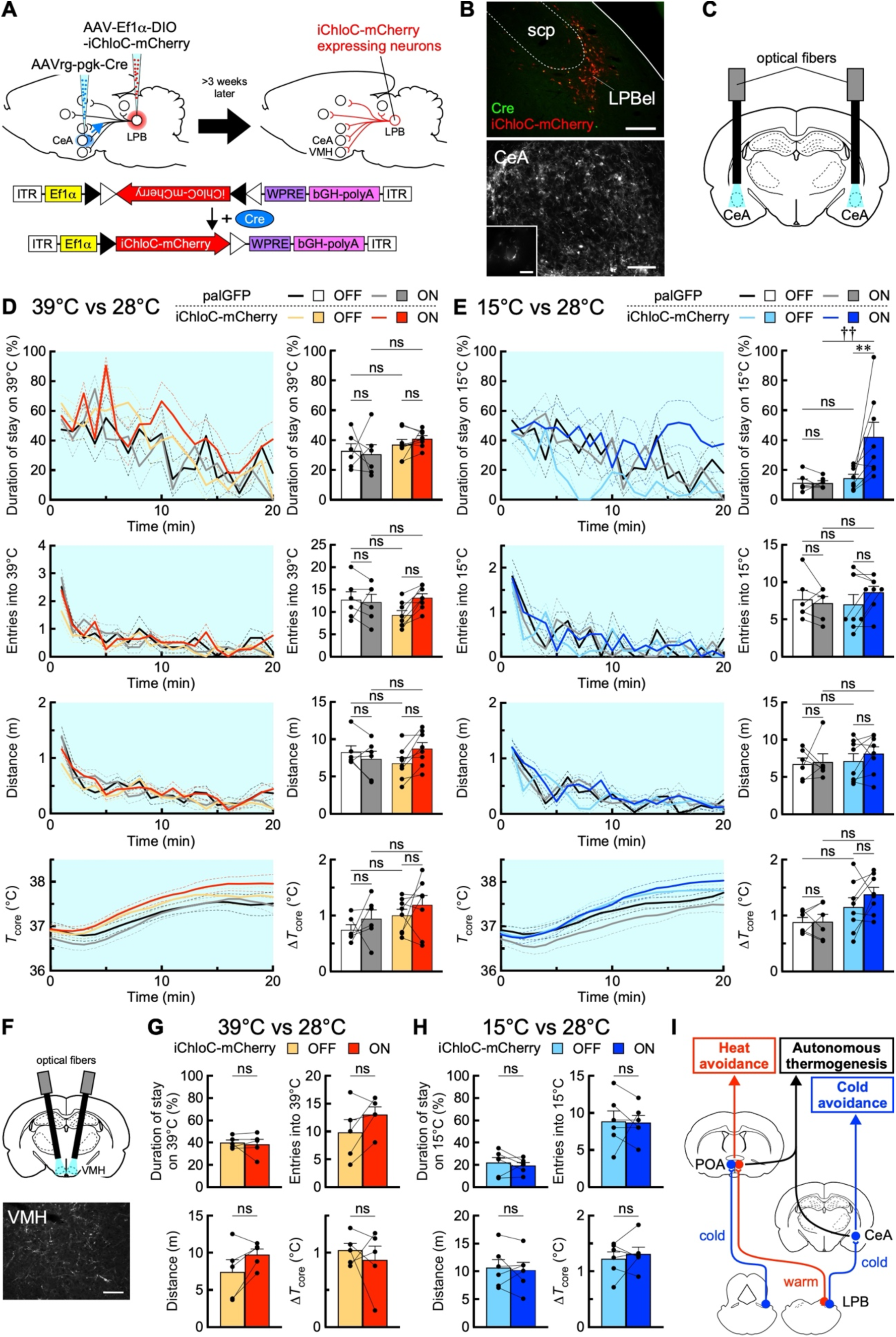
Local optogenetic inhibition of CeA axon terminals of LPB→CeA neurons diminishes cold avoidance. **A**, Selective transduction of LPB→CeA neurons with iChloC-mCherry using AAV and AAVrg. Control rats expressed palGFP instead of iChloC-mCherry. **B**, Cre- and/or iChloC-mCherry-immunoreactive neurons in the LPB (top). Their axon terminals with mCherry immunoreactivity were distributed in the CeA (bottom) and formed basket-like structures (inset). Scale bars, 200 μm (top), 100 μm (bottom), and 10 μm (inset). **C**, CeA axon terminals of LPB→CeA neurons were illuminated. **D and E**, Effects of illumination (light blue) of CeA axon terminals in rats expressing palGFP or iChloC-mCherry in LPB→CeA neurons on heat (D) and cold (E) avoidance. Time-course changes (left, every 1 min) in % duration of stay on 39°C or 15°C plate, times of entry into 39°C or 15°C plate, distance traveled, and *T*_core_. Right graphs show % duration of stay on 39°C or 15°C plate, total times of entry into 39°C or 15°C plate, total distance traveled, and Δ*T*_core_ (difference between value at time 0 and peak) for the 20-min test period, and the data were analyzed by repeated measures one-way ANOVA followed by Bonferroni’s post hoc test [palGFP, *n* = 6, iChloC-mCherry: *n* = 8; (D): duration of stay: *F*_3, 24_ = 1.31, *P* = 0.294; entries: *F*_3, 24_ = 2.00, *P* = 0.141; distance: *F*_3, 24_ = 1.17, *P* = 0.342; Δ*T*_core_: *F*_3, 24_ = 1.73, *P* = 0.187; (E): duration of stay: *F*_3, 24_ = 6.79, *P* = 0.002; entries: *F*_3, 24_ = 0.51, *P* = 0.679; distance: *F*_3, 24_ = 0.43, *P* = 0.736; Δ*T*_core_: *F*_3, 24_ = 3.17, *P* = 0.043). \*\**P* < 0.01; ^††^*P* < 0.01; ns, not significant. **F**, Rats expressing iChloC-mCherry in LPB→CeA neurons received bilateral illumination of the VMH (top). Bottom image shows VMH axon terminals of LPB→CeA neurons expressing iChloC-mCherry. Scale bar, 100 μm. **G and H**, Effects of illumination (light blue) of VMH axon terminals in rats expressing iChloC-mCherry in LPB→CeA neurons on heat (G) and cold (H) avoidance. Graphs show % duration of stay on 39°C or 15°C plate, total times of entry into 39°C or 15°C plate, total distance traveled, and Δ*T*_core_ (difference between value at time 0 and peak) for the 20-min test period, and the data were analyzed by paired *t*-test [(G): *n* = 5; duration of stay, *t*_4_ = 0.43; entries, *t*_4_ = 1.93; distance, *t*_4_ = 2.20; Δ*T*_core_, *t*_4_ = 0.63; (H): *n* = 6; duration of stay, *t*_5_ = 0.71; entries, *t*_5_ = 0.13; distance, *t*_5_ = 0.44; Δ*T*_core_, *t*_5_ = 0.70). ns, not significant. Error bars indicate SEM. **I**, A model for the thermosensory afferent network that drives behavioral and autonomous thermoregulatory responses.

We also photoinhibited VMH collateral axon terminals of LPB→CeA neurons, as the VMH has recently been shown to contain cold-responsive neurons linked to autonomous thermoregulation (Feng et al., 2022). Bilateral photoinhibition of LPB→CeA neuronal axons in the VMH had no effect on heat or cold avoidance or *T*_core_ changes during the TPPTs (Fig. 9, F–H). These results raise the notion that LPB→CeA neurons contribute to cold avoidance via their cold sensory transmission to the CeA, but not via their transmission to the VMH.

## Discussion

This study provides new insights into the importance of LPB-mediated ascending thermosensory pathways in behavioral thermoregulation by demonstrating that synaptic transmission at MnPO axon terminals of LPB→MnPO neurons mediates heat avoidance and that cold sensory transmission to the CeA by LPB→CeA neurons mediates cold avoidance. This study also unexpectedly shows that LPB→CeA neurons, as well as LPB→MnPO neurons, play an important role in autonomous thermoregulation. These findings provide an important framework of the LPB-mediated central thermosensory afferent network that elicits behavioral and autonomous thermoregulatory responses in a coordinated manner to defend thermal homeostasis against environmental thermal challenges. (Fig. 9I)

Our study focused on heat and cold avoidance among thermoregulatory behaviors because it is recruited in the first phase of behavioral thermoregulation in response to changes in ambient temperature. Most previous studies of thermoregulatory behavior have performed operant behavior experiments (Lipton and Hicks, 1968; Carlisle, 1969; Jung et al., 2022). However, because operant thermoregulatory behaviors require a learning process before they are acquired, studying heat and cold avoidance, innate behaviors, is considered more suitable for understanding the fundamental central neural circuit mechanisms of behavioral thermoregulation.

We identified LPB→MnPO neurons and LPB→CeA neurons as segregated subgroups of LPB neurons, which likely belong to the FoxP2- and Lmx1b-expressing groups of LPB neurons, respectively (Huang et al., 2021a, 2021b). Because both groups of LPB neurons are predominantly glutamatergic (Miller et al., 2012; Geerling et al., 2016; Sun et al., 2020), the two groups of LPB projection neurons we focused on probably provide excitatory inputs to the projection sites.

Although both LPB and MnPO have been shown to be pivotal brain sites for autonomous thermoregulation (Morrison and Nakamura, 2019; Nakamura et al., 2022a), whether cutaneous thermosensory monosynaptic inputs from the LPB to the MnPO drive behavioral thermoregulation remains unclear. A recent study showed that optogenetic stimulation of POA axons from *Pdyn*-positive LPB neurons, which are activated by heat exposure, induces postural extension as seen in hot environments (Norris et al., 2021). Because *Pdyn*-positive LPB neurons project to multiple forebrain regions in addition to the POA (Norris et al., 2021), the induction of postural extension by optogenetic stimulation of their POA axons may be due to backfiring to activate collateral axons to another projection site. Moreover, postural extension is elicited by an increase in *T*_core_ rather than *T*_skin_ (Roberts and Martin, 1974), and optogenetic stimulation of POA axons of *Pdyn*-positive LPB neurons had no effect on thermal preference (Norris et al., 2021). In our present study, chemogenetic local presynaptic inhibition of MnPO axon terminals of LPB→MnPO neurons abolished heat avoidance, clearly demonstrating that LPB→MnPO monosynaptic transmission of cutaneous thermosensory information drives heat avoidance. The contribution of *Pdyn*-positive LPB→MnPO neurons to behavioral thermoregulation seems to be limited, and further studies are awaited to identify other LPB neuronal groups responsible for the LPB→MnPO monosynaptic warm sensory transmission for heat avoidance.

Suppression of LPB→MnPO neurons with TeTxLC abolished sympathetic thermogenic and cardiovascular responses to skin cooling and weakened the animals’ ability to defend *T*_core_ in cold and hot environments. These results strongly support the notion that cutaneous cold and warm sensory signaling through the LPB→MnPO pathway elicits feedforward autonomous thermoregulatory responses to environmental thermal challenges (Nakamura and Morrison, 2008, 2010). The cold and warm sensory inputs to the MnPO are considered to alter the intensity of tonic GABAergic inhibitory efferent signaling from prostaglandin EP3 receptor-expressing POA neurons, which controls sympathetic outflow through the DMH and rostral medullary raphe region to thermoregulatory effectors, such as BAT and skin blood vessels, for thermal homeostasis (Nakamura et al., 2022b). Suppression of LPB→MnPO neurons also abolished BAT thermogenesis evoked by a reduction of *T*_core_. Because this thermogenic response was likely elicited by activation of cold-sensitive afferents from body core structures including the abdomen (Gupta et al., 1979), our result suggests that LPB→MnPO neurons transmit afferent signals of temperatures in the peripheral body core as well as the skin. The subgroups of LPB neurons mediating autonomous heat defense responses have been proposed to include those expressing *Pdyn* or *Cck* (Geerling et al., 2016; Yang et al., 2020; Norris et al., 2021), whereas no specific molecular markers have been identified for LPB neurons mediating cold defense.

The upward shift in *T*_core_ caused by suppression of LPB→MnPO neurons at room temperature (Fig. 3A), which probably blocked both cold and warm sensory inputs to the POA, suggests that warm sensory signals to prevent hyperthermia have a more important effect on the POA circuit mechanism for thermal homeostasis than cold sensory signals to prevent hypothermia. Curiously, suppression of LPB→MnPO neurons with TeTxLC had no effect on cold avoidance behavior, suggesting that this pathway is not involved in cold avoidance. However, if LPB→MnPO neurons make a subsidiary contribution to cold avoidance, the effect of blockade of these neurons could be masked by stronger driving signals mediated by other thermosensory pathways, such as the LPB→CeA pathway. Further studies are required to understand the role of LPB→MnPO cold sensory transmission in behavioral thermoregulation.

LPB→CeA neurons have been shown to respond to nociceptive stimuli (Bernard and Besson, 1990; Gauriau and Bernard, 2002). Our present study showed that LPB→CeA neurons are activated in response to cold exposure and contribute to cold avoidance. Because the temperatures used in the cold exposure and cold avoidance test in our study were innocuous, our results indicate that LPB→CeA neurons transmit innocuous cold sensory signals in addition to pain signals. This view is supported by a recent report that LPB-projecting lamina I neurons in the spinal dorsal horn respond polymodally to hot, innocuous cold, and nociceptive stimuli (Chisholm et al., 2021). However, it remains unknown whether the innocuous cold-responsive group of LPB→CeA neurons also responds to other modalities of sensory signals including pain.

Intriguingly, heat avoidance was abolished by chronic suppression of LPB→CeA neurons with TeTxLC, but not by acute photoinhibition of CeA axon terminals of LPB→CeA neurons. Neither did photoinhibition of VMH axon terminals of LPB→CeA neurons affect heat avoidance. These results suggest that heat avoidance requires transmission via axon collaterals of these neurons to a projection site other than the CeA and VMH. Further investigation by photoinhibition of each collateral projection site of LPB→CeA neurons may identify the pathway responsible for mediating heat avoidance. Alternatively, it is possible that a long-term alteration of the neural circuitry by the TeTxLC-mediated chronic suppression of LPB→CeA neurons might have diminished heat avoidance.

Unexpectedly, suppression of LPB→CeA neurons with TeTxLC abolished skin cooling-evoked BAT thermogenesis and cardiovascular responses and also weakened the ability to defend *T*_core_ against cold and hot environmental temperatures. These present results are the first to implicate the amygdala in autonomous responses for thermal homeostasis. Because LPB→CeA neurons projected few collaterals to the POA, it is unlikely that these neurons directly influence the thermoregulatory circuit mechanism in the POA. Distinct from suppression of LPB→MnPO neurons, suppression of LPB→CeA neurons with TeTxLC did not inhibit BAT thermogenesis evoked by a reduction of *T*_core_, suggesting that LPB→CeA neurons do not transmit thermosensory afferent signals from body core structures. In the present study, which focused more on the mechanism of behavioral thermoregulation, we did not test whether optogenetic local inhibition of CeA axon terminals of LPB→CeA neurons alters BAT thermogenesis and other autonomous responses. Further studies are needed to understand how LPB→CeA neurons contribute to autonomous thermoregulation.

The amygdala has been recognized as an important site for the memory of aversive stimuli, such as fear and pain, in order to elicit aversive (avoidance) behavior from the stimuli based on experience (Chiang et al., 2020). However, whether the amygdala stores the memories of innocuous thermal sensations for thermoregulatory behavior has been unknown. Based on our results, it can be hypothesized that cutaneous cold sensory transmission to the CeA through the LPB forms a memory of uncomfortable cold sensation to promote learned cold avoidance behavior. This possibility is supported by our finding that photoinhibition of CeA axon terminals of LPB→CeA neurons significantly diminished cold avoidance only in the last quarter of the test period, by which time a thermosensory memory might be stored in the CeA, whereas chemogenetic blockade of LPB→MnPO synaptic inputs started to inhibit heat avoidance in an earlier phase (Figs. 4D and 9E). The reason why the amygdala stores the memory of cold sensation, but not of warm sensation may be that for small mammals, cold sensation, potentially leading to hypothermia, is more aversive than warm sensation, since moderate heat gain often has a positive value in systemic energy balance. If thermosensory inputs to the CeA generate the unpleasant emotions that underlie innate thermoregulatory behaviors, an important future question is how the unpleasant memories of thermal sensations stored in the amygdala are integrated with the POA-driven commands to develop thermal avoidance behaviors.

## Author contributions

T.Y. and K.N. designed research; T.Y. performed research and analyzed data; N.K. provided AAVs; T.Y., N.K. and K.N. discussed data; T.Y. and K.N. wrote the paper.

## Acknowledgments

We thank Patrick Aebischer and Scott Sternson for sharing their plasmids and Misako Takemoto for technical assistance for AAV production. This study was supported by Moonshot R&D (JPMJMS2023 to K.N.) of the Japan Science and Technology Agency; Grants-in-Aid for Scientific Research (JP22K06470 and JP19K06954 to N.K.; JP23H00398 and JP20H03418 to K.N.) from the Ministry of Education, Culture, Sports, Science and Technology of Japan; and the Japan Agency for Medical Research and Development (JP23wm0525002 to N.K.). T.Y. is supported by the Takeda Science Foundation scholarship.

## References

Almeida, M.C., Steiner, A.A., Branco, L.G.S., Romanovsky, A.A., 2006. Neural substrate of cold-seeking behavior in endotoxin shock. PLoS One 1, e1.

Almeida, M.C., Vizin, R.C.L., Carrettiero, D.C., 2015. Current understanding on the neurophysiology of behavioral thermoregulation. Temperature (Austin) 2, 483–490.

Bernard, J.F., Besson, J.M., 1990. The spino(trigemino)pontoamygdaloid pathway: electrophysiological evidence for an involvement in pain processes. J. Neurophysiol. 63, 473–490.

Carlisle, H.J., 1969. Effect of preoptic and anterior hypothalamic lesions on behavioral thermoregulation in the cold. J. Comp. Physiol. Psychol. 69, 391–402.

Chiang, M.C., Nguyen, E.K., Canto-Bustos, M., Papale, A.E., Oswald, A.-M.M., Ross, S.E., 2020. Divergent Neural Pathways Emanating from the Lateral Parabrachial Nucleus Mediate Distinct Components of the Pain Response. Neuron 106, 927–939.e5.

Chisholm, K.I., Lo Re, L., Polgár, E., Gutierrez-Mecinas, M., Todd, A.J., McMahon, S.B., 2021. Encoding of cutaneous stimuli by lamina I projection neurons. Pain 162, 2405–2417.

Craig, A.D., Bushnell, M.C., Zhang, E.T., Blomqvist, A., 1994. A thalamic nucleus specific for pain and temperature sensation. Nature 372, 770–773.

Craig, A.D., 2002. How do you feel? Interoception: the sense of the physiological condition of the body. Nat. Rev. Neurosci. 3, 655–666.

Feng, C., Wang, Y., Zha, X., Cao, H., Huang, S., Cao, D., Zhang, K., Xie, T., Xu, X., Liang, Z., Zhang, Z., 2022. Cold-sensitive ventromedial hypothalamic neurons control homeostatic thermogenesis and social interaction-associated hyperthermia. Cell Metab. 34, 888–901.e5.

Fulwiler, C.E., Saper, C.B., 1984. Subnuclear organization of the efferent connections of the parabrachial nucleus in the rat. Brain Res. 319, 229–259.

Gauriau, C., Bernard, J.-F., 2002. Pain pathways and parabrachial circuits in the rat. Exp. Physiol. 87, 251–258.

Geerling, J.C., Kim, M., Mahoney, C.E., Abbott, S.B.G., Agostinelli, L.J., Garfield, A.S., Krashes, M.J., Lowell, B.B., Scammell, T.E., 2016. Genetic identity of thermosensory relay neurons in the lateral parabrachial nucleus. Am. J. Physiol. Regul. Integr. Comp. Physiol. 310, R41–54.

Gupta, B.N., Nier, K., Hensel, H., 1979. Cold-sensitive afferents from the abdomen. Pflugers Arch. 380, 203–204.

Han, S., Soleiman, M.T., Soden, M.E., Zweifel, L.S., Palmiter, R.D., 2015. Elucidating an Affective Pain Circuit that Creates a Threat Memory. Cell 162, 363–374.

Hioki, H., Nakamura, H., Ma, Y.-F., Konno, M., Hayakawa, T., Nakamura, K.C., Fujiyama, F., Kaneko, T., 2010. Vesicular glutamate transporter 3-expressing nonserotonergic projection neurons constitute a subregion in the rat midbrain raphe nuclei. J. Comp. Neurol. 518, 668–686.

Huang, D., Grady, F.S., Peltekian, L., Geerling, J.C., 2021a. Efferent projections of Vglut2, Foxp2, and Pdyn parabrachial neurons in mice. J. Comp. Neurol. 529, 657–693.

Huang, D., Grady, F.S., Peltekian, L., Laing, J.J., Geerling, J.C., 2021b. Efferent projections of CGRP/Calca-expressing parabrachial neurons in mice. J. Comp. Neurol. 529, 2911– 2957.

Hylden, J.L., Anton, F., Nahin, R.L., 1989. Spinal lamina I projection neurons in the rat: collateral innervation of parabrachial area and thalamus. Neuroscience 28, 27–37.

Jung, S., Lee, M., Kim, D.-Y., Son, C., Ahn, B.H., Heo, G., Park, J., Kim, M., Park, H.-E., Koo, D.-J., Park, J.H., Lee, J.W., Choe, H.K., Kim, S.-Y., 2022. A forebrain neural substrate for behavioral thermoregulation. Neuron 110, 266–279.e9.

Kataoka, N., Hioki, H., Kaneko, T., Nakamura, K., 2014. Psychological stress activates a dorsomedial hypothalamus-medullary raphe circuit driving brown adipose tissue thermogenesis and hyperthermia. Cell Metab. 20, 346–358.

Kataoka, N., Shima, Y., Nakajima, K., Nakamura, K., 2020. A central master driver of psychosocial stress responses in the rat. Science 367, 1105–1112.

Li, J., Xiong, K., Pang, Y., Dong, Y., Kaneko, T., Mizuno, N., 2006. Medullary dorsal horn neurons providing axons to both the parabrachial nucleus and thalamus. J. Comp. Neurol. 498, 539–551.

Link, E., Edelmann, L., Chou, J.H., Binz, T., Yamasaki, S., Eisel, U., Baumert, M., Südhof, T.C., Niemann, H., Jahn, R., 1992. Tetanus toxin action: inhibition of neurotransmitter release linked to synaptobrevin proteolysis. Biochem. Biophys. Res. Commun. 189, 1017–1023.

Lipton, J.M., Hicks, S.P., 1968. Effects of Preoptic Lesions on Heat-escape Responding and Colonic Temperature in the Rat’. Pergamon Press.

Miller, R.L., Knuepfer, M.M., Wang, M.H., Denny, G.O., Gray, P.A., Loewy, A.D., 2012. Fos-activation of FoxP2 and Lmx1b neurons in the parabrachial nucleus evoked by hypotension and hypertension in conscious rats. Neuroscience 218, 110–125.

Morrison, S.F., Nakamura, K., 2019. Central Mechanisms for Thermoregulation. Annu. Rev. Physiol. 81, 285–308.

Nakamura, K., Kaneko, T., Yamashita, Y., Hasegawa, H., Katoh, H., Negishi, M., 2000. Immunohistochemical Localization of Prostaglandin EP3 Receptor in the Rat Nervous System.

Nakamura, K., Matsumura, K., Hübschle, T., Nakamura, Y., Hioki, H., Fujiyama, F., Boldogköi, Z., König, M., Thiel, H.-J., Gerstberger, R., Kobayashi, S., Kaneko, T., 2004. Identification of sympathetic premotor neurons in medullary raphe regions mediating fever and other thermoregulatory functions. J. Neurosci. 24, 5370–5380.

Nakamura, K., Morrison, S.F., 2007. Central efferent pathways mediating skin cooling-evoked sympathetic thermogenesis in brown adipose tissue. Am. J. Physiol. Regul. Integr. Comp. Physiol. 292, R127–36.

Nakamura, K., Morrison, S.F., 2008. Preoptic mechanism for cold-defensive responses to skin cooling. J. Physiol. 586, 2611–2620.

Nakamura, K., Morrison, S.F., 2010. A thermosensory pathway mediating heat-defense responses. Proc. Natl. Acad. Sci. U. S. A. 107, 8848–8853.

Nakamura, K., Morrison, S.F., 2011. Central efferent pathways for cold-defensive and febrile shivering. J. Physiol. 589, 3641–3658.

Nakamura, K., Nakamura, Y., Kataoka, N., 2022a. A hypothalamomedullary network for physiological responses to environmental stresses. Nat. Rev. Neurosci. 23, 35–52.

Nakamura, Y., Yahiro, T., Fukushima, A., Kataoka, N., Hioki, H., Nakamura, K., 2022b. Prostaglandin EP3 receptor-expressing preoptic neurons bidirectionally control body temperature via tonic GABAergic signaling. Sci Adv 8, eadd5463.

Norris, A.J., Shaker, J.R., Cone, A.L., Ndiokho, I.B., Bruchas, M.R., 2021. Parabrachial opioidergic projections to preoptic hypothalamus mediate behavioral and physiological thermal defenses. Elife 10, 1–29.

Roberts, W.W., Martin, J.R., 1974. Peripheral thermoreceptor control of thermoregulatory responses of the rat. J. Comp. Physiol. Psychol. 87, 1109–1118.

Saper, C.B., Loewy, A.D., 1980. Efferent connections of the parabrachial nucleus in the rat. Brain Res. 197, 291–317.

Satinoff, E., Rutstein, J., 1970. Behavioral thermoregulation in rats with anterior hypothalamic lesions. J. Comp. Physiol. Psychol. 71, 77–82.

Sato, M., Ito, M., Nagase, M., Sugimura, Y.K., Takahashi, Y., Watabe, A.M., Kato, F., 2015. The lateral parabrachial nucleus is actively involved in the acquisition of fear memory in mice. Mol. Brain 8, 22.

Schiavo, G., Benfenati, F., Poulain, B., Rossetto, O., Polverino de Laureto, P., DasGupta, B.R., Montecucco, C., 1992. Tetanus and botulinum-B neurotoxins block neurotransmitter release by proteolytic cleavage of synaptobrevin. Nature 359, 832–835.

Stachniak, T.J., Ghosh, A., Sternson, S.M., 2014. Chemogenetic synaptic silencing of neural circuits localizes a hypothalamus→midbrain pathway for feeding behavior. Neuron 82, 797‒808.

Sun, L., Liu, R., Guo, F., Wen, M.-Q., Ma, X.-L., Li, K.-Y., Sun, H., Xu, C.-L., Li, Y.-Y., Wu, M.-Y., Zhu, Z.-G., Li, X.-J., Yu, Y.-Q., Chen, Z., Li, X.-Y., Duan, S., 2020. Parabrachial nucleus circuit governs neuropathic pain-like behavior. Nat. Commun. 11, 5974.

Takahashi, M., Ishida, Y., Kataoka, N., Nakamura, K., Hioki, H., 2021. Efficient Labeling of Neurons and Identification of Postsynaptic Sites Using Adeno-Associated Virus Vector, in: Lujan, R., Ciruela, F. (Eds.), Receptor and Ion Channel Detection in the Brain. Springer US, New York, NY, pp. 323–341.

Tamamaki, N., Nakamura, K., Furuta, T., Asamoto, K., Kaneko, T., 2000. Neurons in Golgi-stain-like images revealed by GFP-adenovirus infection in vivo. Neurosci. Res. 38, 231–236.

Tan, C.L., Cooke, E.K., Leib, D.E., Lin, Y.-C., Daly, G.E., Zimmerman, C.A., Knight, Z.A., 2016. Warm-Sensitive Neurons that Control Body Temperature. Cell 167, 47–59.e15.

Wietek, J., Beltramo, R., Scanziani, M., Hegemann, P., Oertner, T.G., Wiegert, J.S., 2015. An improved chloride-conducting channelrhodopsin for light-induced inhibition of neuronal activity in vivo. Sci. Rep. 5, 14807.

Yahiro, T., Kataoka, N., Nakamura, Y., Nakamura, K., 2017. The lateral parabrachial nucleus, but not the thalamus, mediates thermosensory pathways for behavioural thermoregulation. Sci. Rep. 7, 5031.

Yang, W.Z., Du, X., Zhang, W., Gao, C., Xie, H., Xiao, Y., Jia, X., Liu, J., Xu, J., Fu, Xin, Tu, H., Fu, Xiaoyu, Ni, X., He, M., Yang, J., Wang, H., Yang, H., Xu, X.-H., Shen, W.L., 2020. Parabrachial neuron types categorically encode thermoregulation variables during heat defense. Sci Adv 6, 1–17.

Yu, S., Qualls-Creekmore, E., Rezai-Zadeh, K., Jiang, Y., Berthoud, H.-R., Morrison, C.D., Derbenev, A.V., Zsombok, A., Münzberg, H., 2016. Glutamatergic Preoptic Area Neurons That Express Leptin Receptors Drive Temperature-Dependent Body Weight Homeostasis. J. Neurosci. 36, 5034–5046.

